# High homozygosity of inversions in sunflower species largely averts accumulation of deleterious mutations

**DOI:** 10.1101/2022.01.06.475294

**Authors:** Kaichi Huang, Kate L. Ostevik, Cassandra Elphinstone, Marco Todesco, Natalia Bercovich, Gregory L. Owens, Loren H. Rieseberg

## Abstract

Recombination is critical both for accelerating adaptation and for the purging of deleterious mutations. Chromosomal inversions can act as recombination modifiers that suppress local recombination and, thus, are predicted to accumulate such mutations. In this study, we investigated patterns of recombination, transposable element abundance and coding sequence evolution across the genomes of 1,445 individuals from three sunflower species, as well as within nine inversions segregating within species. We also analyzed the effects of inversion genotypes on 87 phenotypic traits to test for overdominance. We found significant negative correlations of long terminal repeat retrotransposon abundance and deleterious mutations with recombination rates across the genome in all three species. However, we failed to detect an increase in these features in the inversions, except for a modest increase in the proportion of stop codon mutations in several very large or rare inversions. Moreover, there was little evidence of phenotypic overdominance in inversion heterozygotes, consistent with observations of minimal deleterious load. On the other hand, significantly greater load was observed for inversions in populations polymorphic for a given inversion compared to populations monomorphic for one of the arrangements, suggesting that the local state of inversion polymorphism affects deleterious load. These seemingly contradictory results can be explained by the geographic structuring and consequent excess homozygosity of inversions in wild sunflowers. Inversions contributing to local adaptation often exhibit geographic structure; such inversions represent ideal recombination modifiers, acting to facilitate adaptive divergence with gene flow, while largely averting the accumulation of deleterious mutations due to recombination suppression.

## INTRODUCTION

Chromosomal inversions are thought to play an important role in adaptation and speciation. In one class of models, inversions contribute to adaptive evolution by reducing gene flow over large genomic regions, while maintaining advantageous epistatic interactions and/or favorable genotypic combinations at loci affecting adaptation to different environments (Charlesworth and Charlesworth 1973; Kirkpatrick and Barton 2006; Feder et al. 2011; Charlesworth and Barton 2018). Other models suggest that inversions can facilitate speciation by increasing effective linkage between genes that contribute to local adaptation and those causing assortative mating (Trickett and Butlin 1994; Ortiz-Barrientos et al. 2016) or by enabling the accumulation of intrinsic genetic incompatibilities (Noor et al. 2001; Navarro and Barton, 2003).

These recombination suppression models offer a means for resolving the widely-recognized antagonism between divergent natural selection and recombination (Felsenstein 1981). However, genetic recombination between homologous chromosomes is essential for removing deleterious mutations from the genome (Carvalho 2003; Keightley and Otto 2006). In genomic regions of low recombination, the efficiency of selection in eliminating slightly deleterious mutations is expected to weaken because deleterious mutations might hitchhike with genes that are under positive selection (i.e., Hill-Robertson effects; Hill and Robertson 1966; Morrell et al. 2011). Population genomic analyses of *Drosophila* and other animal species have provided evidence for this evolutionary advantage of genetic recombination (Bachtrog 2003; Kaiser and Charlesworth 2009; Wang et al. 2013; Yoshida et al. 2020). Likewise, studies of a number of crops have reported a negative correlation between mutational load and recombination rate (Lu et al. 2006; Renaut and Rieseberg 2015; Lozano et al. 2021), consistent with a critical role for recombination in plants as well.

In inversions, effective recombination between the arrangements is reduced through selection against recombinant gametes in heterokaryotypes or via crossover interference mechanisms (Fuller et al. 2019; Huang and Rieseberg 2020). Such suppression of recombination promotes independent evolution of each arrangement and may result in a decrease in effective population size within each arrangement. Therefore, the efficacy of purifying selection is expected to be lower in inversions, making them more vulnerable to the accumulation of deleterious mutations when compared to the rest of the genome (Faria et al. 2019; Berdan et al. 2021). While this effect is expected to be stronger in rare arrangements (Montgomery et al. 1987; Navarro, et al. 2000), theoretical work indicates that excess deleterious load can accumulate in both arrangements (Berdan et al. 2021).

Masking of recessive deleterious mutations that are restricted to only one inversion arrangement may result in higher fitness in heterokaryotypes than in homokaryotypes. Such associative overdominance favors intermediate-to-large inversions that persist as balanced polymorphisms (Ohta 1971; Connallon and Olito 2021; Faria et al. 2019; Berdan et al. 2021). While a similar inversion length distribution is predicted under the local adaptation with gene flow model, an inversion arrangement might spread to fixation by selection in some local populations while being absent or lost in others. Inversions in this case should display higher homozygosity than those established via associative overdominance and be less prone to the accumulation of deleterious alleles due to the large local effective population size of each arrangement (Faria et al. 2019).

Associative overdominance has been showed to be an important force for maintaining polymorphism in a number of inversions in Insecta, such as seaweed flies *Coelopa frigida* (Butlin and Day 1985) and *Heliconius* butterflies (Jay et al. 2021). However, we are unaware of comparable studies of plant inversions, many of which established in the scenario of local adaptation with gene flow (Lowry and Willis 2010; Lee et al. 2017; Coughlan and Willis 2019; Huang et al. 2020).

In population genomic studies, deleterious load is typically detected by the study of protein coding genes. For example, within a species one can compare the average number of nonsynonymous differences per nonsynonymous site (referred to as π_N_) to the average number of synonymous differences per synonymous site (π_S_) to estimate the effect on selection on protein sequences (Hollister et al. 2015; Hahn 2019). An increase of this ratio usually indicates an increase in load of deleterious mutations within the population. In practice, researchers also use the ratio of the diversity at zero-fold degenerate sites and the diversity at four-fold degenerate sites (π_0_/π_4_) as an approximation (Marsden et al. 2016; Yang et al. 2018; Chen et al. 2020). Zero-fold degenerate sites are sites where any nucleotide substitution will result in amino acid substitution, which is an approximation of nonsynonymous sites, while four-fold degenerate sites are those where four possible nucleotides will result in the same amino acid. In addition, nonsense mutations that cause premature termination of protein synthesis are normally highly deleterious and subjective to purifying selection (Chu and Wei 2019). Therefore, an increase in the proportion of nonsense mutations relative to total number of nonsynonymous mutations would also indicate increasing deleterious load (Renaut and Rieseberg 2015).

Transposable elements (TEs) represent another potentially abundant source of deleterious mutations. Not only are they the largest component of plant and animal genomes (typically > 50%), but they also contribute disproportionately to genomic variation within and between species. TE insertions, for example, often produce detrimental effects on fitness because they may disrupt gene sequences and/or alter gene expression and function (Lisch 2013). In addition, ectopic recombination between non-homologous element copies can produce deleterious chromosome rearrangements (Montgomery et al. 1987; Langley et al. 1988). Therefore, it is predicted that TEs will be more abundant in chromosome regions that have reduced levels of recombination, both because selection acting against deleterious mutations is weaker in regions of low recombination and because TEs are less likely to be involved in unequal recombination between elements located on homologous chromosomes in those regions (Langley et al. 1988; Ma and Bennetzen 2006; Kent et al. 2017). This effect of reduced recombination in TE accumulation is supported by a number of studies in *Drosophila* showing that TE abundance is strongly associated with the local recombination rate (Langley et al. 1988; Bartolome et al. 2002). In plants, recombination rates are significantly associated with long terminal repeat retrotransposon (LTR-RT) abundance in rice (Tian et al. 2009), soybean (Tian et al. 2012), and wheat (Daron et al. 2014), as well as the frequency of different classes of TEs in *Eucalyptus* (Gion et al. 2016). However, examination of the genome of *Arabidopsis thaliana* found that TE abundance did not correlate with recombination rate, perhaps because inbreeding reduces the effects of recombination rate variation (Wright et al. 2003).

Because chromosomal inversions can reduce recombination, it has also been proposed that TEs should accumulate in inverted regions (Montgomery et al. 1987; Sniegowski and Charlesworth 1994). Such an increase in TE copy number within inversions has been reported in *Drosophila* (Eanes et al. 1992; Sniegowski and Charlesworth 1994) as well as in *Heliconius* (Jay et al. 2021). However, we are unaware of such reports in plants.

Sunflowers are an excellent system to study the effects of inversions on the accumulation of deleterious mutations. First, large non-recombining haplotype blocks (haploblocks) have been found to play a critical role in ecotype formation in wild sunflowers (Todesco et al. 2020): seven of them underlie the formation of the *texanus* ecotype of common sunflower, *Helianthus annuus*; four gave rise to an early-flowering ecotype of the silverleaf sunflower, *H. argophyllus*; and seven were found to be associated with adaptive traits (e.g. seed size) in well-characterized dune ecotypes of prairie sunflower, *H. petiolaris*. Many of these haploblocks were found to be linked to polymorphic chromosomal inversions (Todesco et al.

2020). Second, an increase in deleterious mutations, such as nonsynonymous substitutions and alternative stop codons, has previously been reported in regions of reduced recombination in cultivated sunflower (Renaut and Rieseberg 2015), suggesting that it should be possible to detect an effect of inversions on protein evolution in sunflower species, if it exists. Third, more than three quarters of the sunflower genome is comprised of TEs, of which 77% are LTR-RTs (Staton et al. 2012; Badouin et al. 2017). Sunflower LTR-RTs are predominantly of recent origin and exhibit transcriptional activity among wild species (Cavallini et al. 2010; Kawakami et al. 2011; Renaut et al. 2014). However, whether and how the accumulation pattern of TEs is shaped by inversions has not previously been explored in sunflowers.

In this study, we investigated the distribution of TEs, specifically LTR-RTs, and coding sequence evolution across the sunflower genome, as well as within nine inversions identified in a previous study in multiple wild sunflower species (Todesco et al. 2020; table 1). We focused on the impact of recombination and inversions on TE density and the accumulation of deleterious mutations in protein coding genes, quantified as elevated nonsynonymous diversity and increased incidence of nonsense mutations. We further looked for a signal of elevated genetic load in inversions by testing for overdominance of phenotypic traits. In addition, we mapped the location of centromeres in the sunflower genome both to assess their impact on recombination rates and to determine if inversions included centromeres (pericentric) or not (paracentric). We showed that variation in recombination rate along the sunflower genome plays a crucial role in the accumulation of TEs and in modulating the efficacy of selection in removing deleterious mutations. However, we found that inversions established under the local adaptation scenario (7 paracentric and 2 pericentric), which displayed higher than expected homozygosity across populations, had little impact on deleterious load.

**Table 1.**
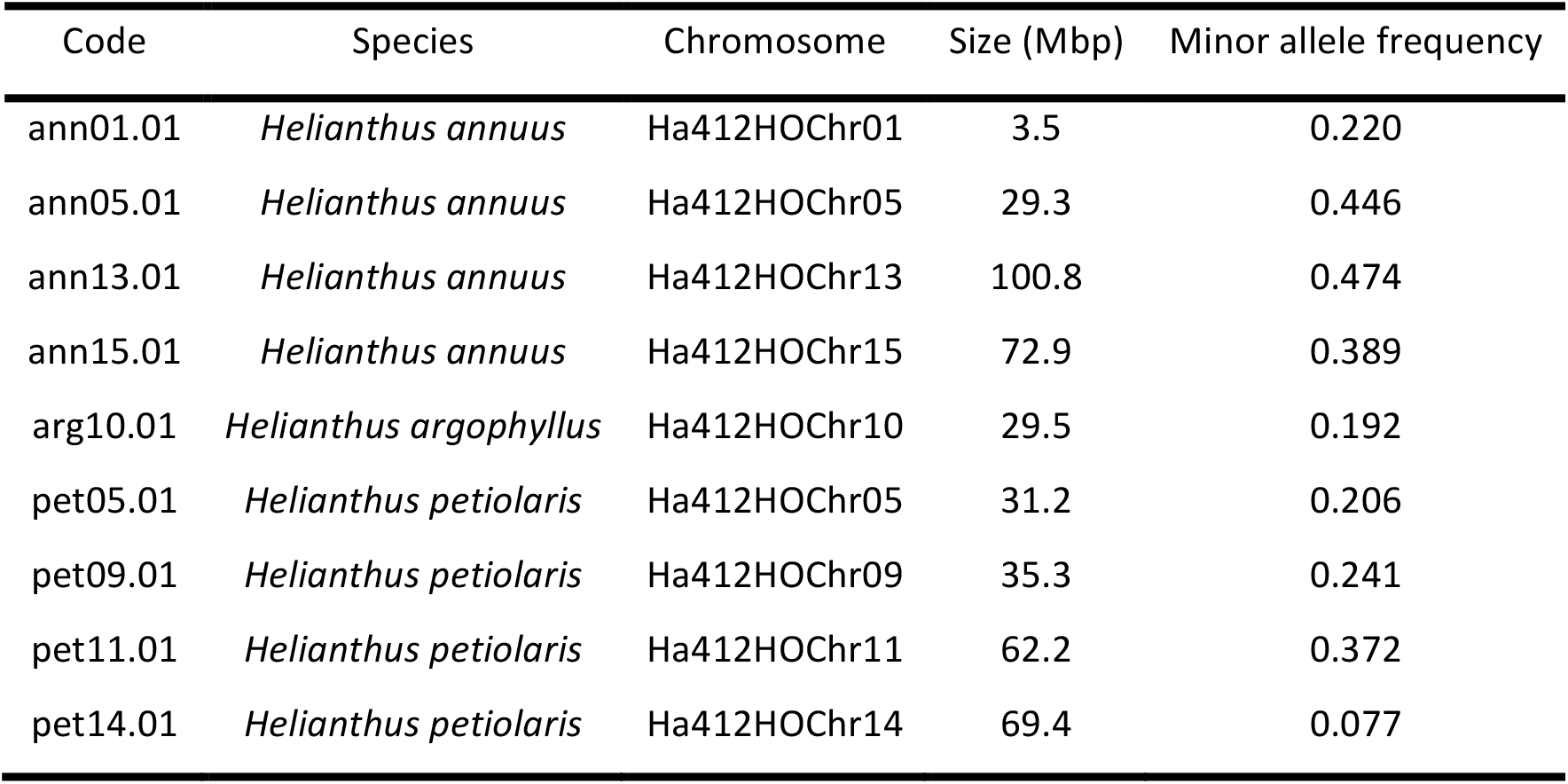
Codes and basic features of the inversions in the analyses.

## RESULTS

### SNP dataset

In this study, we used the whole-genome sequencing data of three annual sunflower species (*H. annuus, H. argophyllus, H. petiolaris*) from a previous study (Todesco et al. 2020) and generated whole-genome sequencing data for an additional 38 samples from two dune systems in *H. petiolaris*. We conducted variant calling on a new high-quality reference genome for *H. annuus* (Ha412HOv2.0), which used Hi-C (Marie-Nelly et al. 2014) for contig and scaffold ordering and was shown to have improved quality (Todesco et al. 2020). A total of 71 populations and 719 samples for *H. annuus*, 30 populations and 299 samples for *H. argophyllus* and 43 populations and 427 samples for *H. petiolaris* were included in this study (see Materials and Methods). After calling variants and filtering, we obtained 15,452,562, 8,706,003, and 8,797,015 bi-allelic SNPs for *H. annuus, H. argophyllus* and *H. petiolaris*, respectively. This is approximately double the number SNPs included in our previous datasets, despite application of similar filtering standards (Todesco et al. 2020).

### Genomic recombination rate and centromeres

To explore the effect of recombination on molecular evolution across the genome, we made use an integrated genetic map for cultivated sunflower (Todesco et al. 2020) and remapped the markers to our current reference genome. This map integrated genetic maps from seven crosses to produce an average recombination rate for the species. Analysis of patterns of recombination rate variation across the genome of Ha412HOv2.0 using the integrated genetic map showed that most of the chromosomes have large regions of low recombination around central regions and high recombination toward the distal ends. Some chromosomes, such as Ha412HOChr06, have regions of low recombination near one terminal end (fig. 1). Since the recombination rate is relatively constant among annual sunflower species (Barb et al. 2014), we used the recombination map for *H. annuus* to estimate patterns of recombination in all three species in our study.

**Fig 1.**
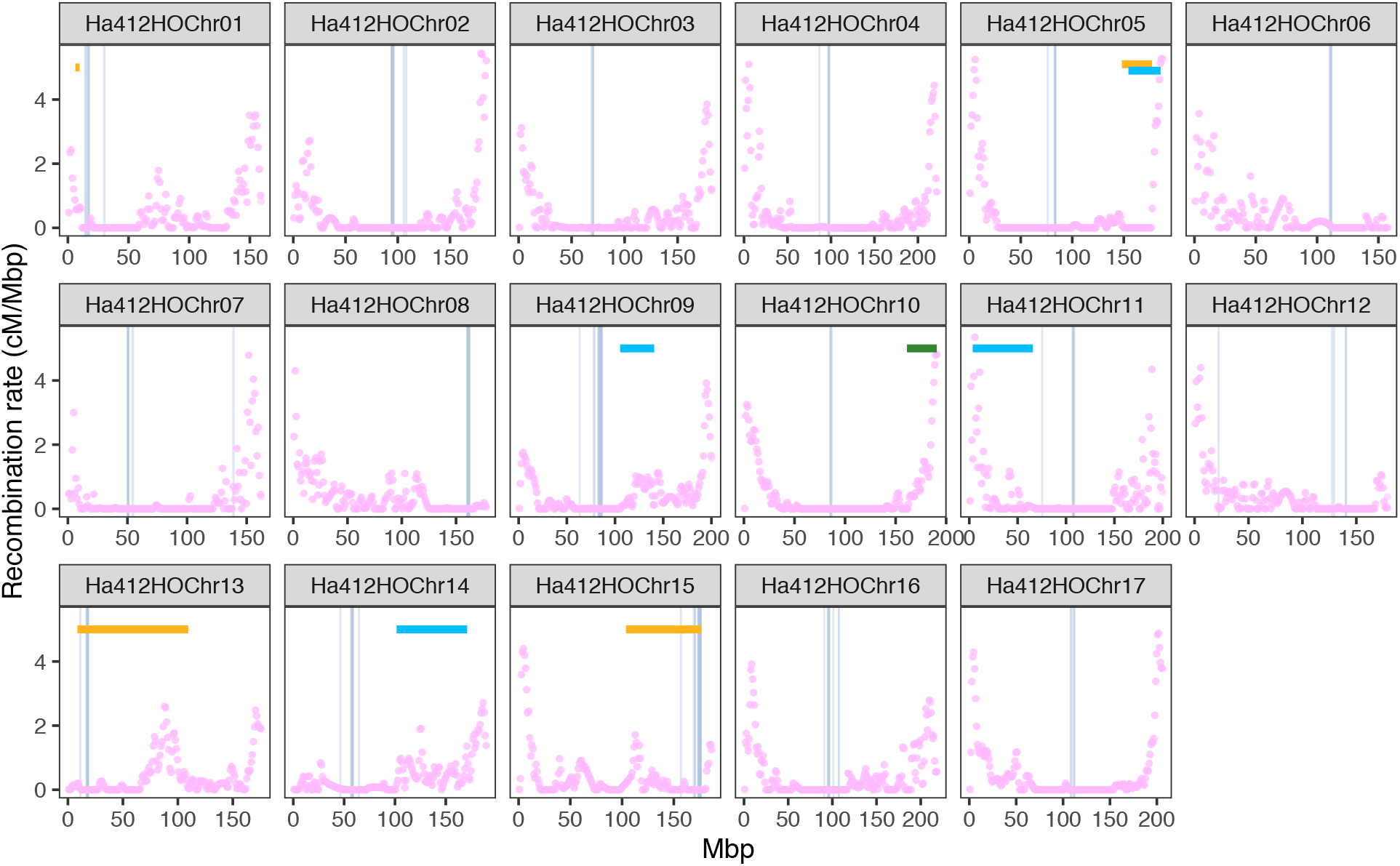
Recombination rate across the genome of Ha412HOv2.0. The recombination rate was calculated for 500-kbp windows based on an integrated genetic map of cultivated sunflower (*Helianthus annuus*). Positions of the centromere-specific sequence LC075745 are indicated by grey vertical bars. Positions of the inversions used in this study are indicated by horizontal bars (orange= *H. annuus*, green= *H. argophyllus*, light blue= *H. petiolaris*).

We mapped four sequences previously found to be targeted by sunflower centromere-specific histone H3 (LC075744-LC075747; Nagaki et al. 2015) to the reference genome to determine the position of centromeres. Among the four sequences, the LINE-like sequence LC075745 was mapped to a single centralized region on most chromosomes, except for chromosomes 7 and 12 where the sequence was also mapped to another small region (fig. 1; supplementary fig. S1, Supplementary Material online). The other three tandem repeat/retrotransposon sequences showed less well-defined patterns. Most of the hits of LC075744 were located on un-anchored scaffolds. Other hits were found on 7 out of 17 chromosomes, the majority of which appeared in the vicinity of corresponding hits for LC075745. Sequence LC075746 was mapped to wider regions across the chromosomes but mostly appeared densely around the region identified by LC075745. Sequence LC075747 was found throughout the genome (supplementary fig. S1, Supplementary Material online). From this, we infer that the centromere locations were characterized by the regions delineated by LC075745. As expected, all the centromeres were located within a region of extremely low recombination, although the centromere of chromosome 1 was on the edge of the region; chromosomes 6, 8 and 13 are acrocentric, which fit the distribution of recombination rate on these chromosomes (fig. 1). Among the inversions, ann13.01 on chromosome 13 and ann15.01 on chromosome 15 are pericentric while the others are paracentric (fig. 1).

### Inversions suppress recombination between arrangements

Using the newly generated SNP dataset, we examined patterns of linkage disequilibrium (LD) across our samples and sequence divergence between arrangements to confirm the effect of recombination suppression for the inversions in sunflowers. When we compared LD between samples from populations that are polymorphic for the inversion and those from populations that contain only the homozygous genotype for the major arrangement, all inversions were characterized by a region of high LD among samples from polymorphic populations, while this pattern was not found among samples from populations with homozygous genotypes, except that a small region of high LD persisted in the centromeric regions in paracentric inversions ann13.01 and ann15.01 (supplementary fig. S2, Supplementary Material online). These results are consistent with the role of inversions in altering recombination in heterozygotes, while recombination in homozygotes remains unaffected.

In most of the inversions, divergence between arrangements is consistently high across the whole region relative to the low *F*_ST_ outside the rearranged regions (supplementary fig. S3, Supplementary Material online), except for the abrupt gaps in the middle of pet05.10 and pet09.01. These result from the structural differences between *H. petiolaris* and the reference genome, which is for cultivated sunflower (*H. annuus* var. *macrocarpus*; Ostevik et al. 2020). In the two largest and pericentric inversions, ann13.01 and ann15.01, *F*_ST_ is more heterogeneous with a gradual decline in the middle of the inverted region, consistent with increased double recombination events and gene conversion between arrangements away from the breakpoints.

### Transposable element abundance

To explore the effect of inversions on TE abundance, we made use of the three reference genomes of cultivated sunflowers (Ha412HOv2.0, XRQv2, PSC8) that have been found to contain different arrangements of two inversions found in the wild populations (Todesco et al. 2020): Ha412HOv2.0 and XRQv2 have the same arrangement for the inversion on chromosome 1 (ann01.01), while PSC8 has the alternate; XRQv2 and PSC8 possess the same arrangement for the inversion on chromosome 5 (ann05.01), while Ha412HOv2.0 has the alternate. To generate comparable datasets, we conducted de-novo TE annotation for the three reference genomes using the program EDTA, which is a sophisticated pipeline that creates raw TE libraries using various structure-based programs and filters out false discoveries in raw TE candidates to generate a high-quality non-redundant TE library (Ou et al. 2016). Using the EDTA pipeline to annotate the three genome assemblies, we confirmed that the genomes of cultivated sunflower are composed of a high proportion of TEs (83.02% in Ha412HOv2.0, 83.33% in XRQv2, 83.21% in PSC8), with 71-72% of these genomes being LTR-RTs (71.50% in Ha412HOv2.0, 71.20% in XRQv2, 71.29% in PSC8). In agreement with previous studies of the cultivated sunflower genome (Staton et al. 2012), there is a major bias in TE composition towards *Gypsy* (Ha412HOv2.0 42.04%, XRQv2 41.93%, PSC8 41.63%) and *Copia* elements (Ha412HOv2.0 11.12%, XRQv2 12.04%, PSC8 12.52%). Class II TEs (DNA transposons) were much lower in abundance relative to LTR-RTs, comprising about 12% of each genome (Ha412HOv2.0 11.50%, XRQv2 12.14%, PSC8 11.92%).

The genomic distributions of LTR-RTs in the three assemblies are similar to those previously reported for the first reference genome for cultivated sunflower (Badouin et al. 2017; supplementary fig. S4, Supplementary Material online). We found a strong negative correlation between density of LTR-RTs and recombination rates (p < 2.2×10^−16^; fig. 2a). The results were similar when considering only *Gypsy* or *Copia* elements (not shown). Our integrated genetic map has not been aligned to the genomes of XRQv2 and PSC8, but the similar distributions of LTR-RTs across the three genomes imply a comparably strong negative correlation between recombination rate and TE abundance.

**Fig 2.**
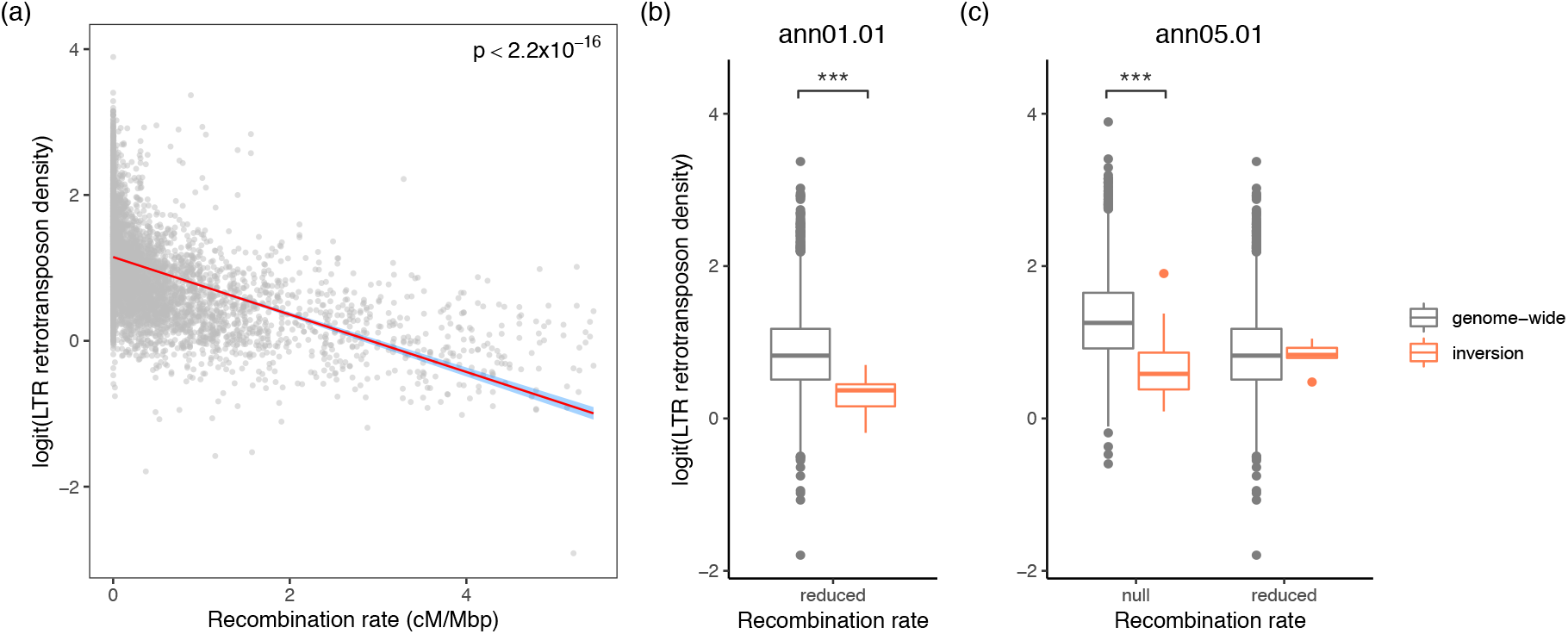
Recombination rate and long terminal repeat (LTR) retrotransposon density. (a) Genome-wide correlation of LTR retrotransposon density and recombination rate in Ha412HOv2.0. The red line denotes the best-fit linear regression line with 95% confidence intervals shaded in blue. (b) Comparison of LTR retrotransposon density within inversion ann01.01 to genome-wide. (c) Comparison of the LTR retrotransposon density within inversion ann05.01 to genome-wide. Windows in each recombination rate category (reduced: 0.01-2 cM/Mbp, null: <0.01 cM/Mbp) were compared separately. Asterisks denote significance in independent t-test: ***p<0.001.

We divided 500 kbp windows of the genome into three different categories based on the recombination rate (“high”: >2 cM/Mbp, “reduced”: 0.01-2 cM/Mbp, “null”: <0.01 cM/Mbp), and compared windows of each category in an inversion to those of the same category from across the genome, to control for the effect of genomic context on TE density (Bartolome et al. 2002). We note that although neighboring genomic windows are not completely independent, linkage typically declines within about 10 kbp in sunflowers (Todesco et al. 2020). For the two inversions that are polymorphic between the reference genomes, the recombination rate of most windows in ann01.01 fell within the “reduced” category, whereas for ann05.01, both “reduced” and “null” windows were found. Recall that these recombination rates are derived from genetic mapping populations that are homozygous for the inversions. Thus, this is the background recombination rate for these windows rather than the recombination rate in the inversion heterozygotes. When compared to windows of a similar recombination rate across the genome, ann01.01 exhibited significantly lower density of LTR-RTs (p = 9.17×10^−4^; fig. 2b). Likewise, for ann05.01, windows with a “null” recombination rate contained fewer LTR-RTs than windows with a similar recombination rate elsewhere in the genome (p = 6.62×10^−22^), but no significant difference was found between windows of the “reduced” recombination rate category across the genome and those within the inversion (p = 0.557; fig. 2c). LTR-RT densities in different inversion arrangements from multiple genome assemblies did not show visible differences and were all lower than the genomic average (supplementary fig. S5, Supplementary Material online).

### Protein evolution

We selected 20 samples from *H. annuus, H. argophyllus* and *H. petiolaris*, respectively, and summarized zero-fold and four-fold diversity across the whole genome to estimate the genetic load of these samples. Across the three species, π_0_/π_4_ shows a negative correlation with levels of neutral heterozygosity (mean of π_0_ and π_4_; supplementary fig. S6, Supplementary Material online). *Helianthus argophyllus* samples have the lowest levels of neutral diversity among the species. However, they show disproportionately higher levels of diversity at zero-fold sites (supplementary fig. S6, Supplementary Material online). *Helianthus annuus* has similar but slightly higher levels of neutral diversity and lower levels of π_0_/π_4_ compared to *H. petiolaris*. To test the significance of this inverse relationship while controlling for spurious signal caused by mathematical correlation between π_0_/π_4_ and genetic diversity, both π_0_ and π_4_ were scaled by dividing by the standard deviation of each, and the association between the average of the scaled values and the ratio between them were then tested (Irwin, 2018). This resulted in a statistically significant inverse relationship (Pearson’s r = -0.7116, p = 1.848×10^−10^).

In all three species, the ratio of nonsynonymous and synonymous mutations as estimated by window-based π_0_/π_4_ is similar to that seen in other taxa (Yang and Gaut 2011) with a small proportion of the values greater than 2 (supplementary fig. S7, Supplementary Material online). Mean π_0_/π_4_ ranges from 0.3-0.5, with the highest value found in *H. argophyllus* and the lowest in *H. petiolaris* (*H. annuus*: 0.3787; *H. argophyllus*: 0.4248; *H. petiolaris*: 0.3124). Across the genome, π_0_/π_4_ is negatively correlated with recombination rate in all three species, but the correlation is weak in *H. petiolaris* (*H. annuus*: Pearson’s r = -0.1162, p < 2.2×10^−16^; *H. argophyllus*: Pearson’s r = -0.1752, p < 2.2×10^−16^; *H. petiolaris*: Pearson’s r = - 0.03836, p = 0.00448; fig. 3a-c).

**Fig 3.**
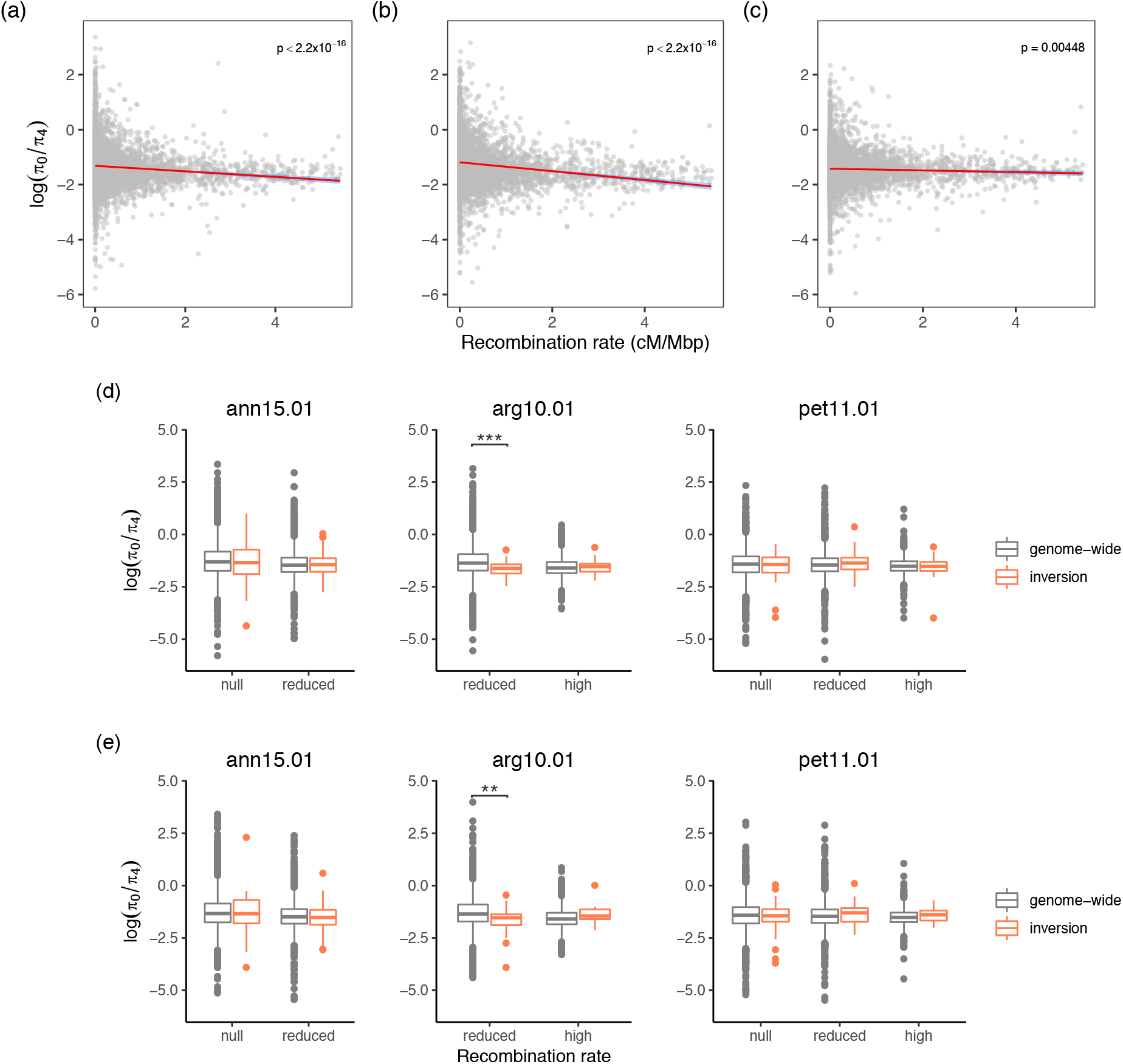
Recombination rate and π_0_/π_4_. Genome-wide correlation of π_0_/π_4_ and recombination rate in (a) *Helianthus annuus*, (b) *H. argophyllus* and (c) *H. petiolaris*. The red lines denote the best-fit linear regression line with the 95% confidence intervals shaded in blue. The values within inversions were compared to genome-wide regions in the same recombination rate category using (d) all samples or (e) samples homozygous for the rarer arrangement. Windows of each recombination rate category were compared separately. Asterisks denote significance in independent t-test: **0.01>p>0.001; ***p<0.001. Results for all inversions are presented in supplementary fig. S8 and S9, Supplementary Material online.

We compared π_0_/π_4_ for the inversions to genome-wide measures in a similar way to that in the above analysis of TE density. Contrary to what was expected, most of the inversions that we examined showed no difference in deleterious load (p > 0.1) when compared to the genomic background in each recombination rate category. In the few comparisons where significant (p < 0.01) results were obtained, such as for windows with “reduced” recombination rate in inversion arg10.01, the inversions have lower π_0_/π_4_ values, indicating fewer nonsynonymous substitutions in those windows than the genome-wide average for regions with similar recombination rates (fig. 3d; supplementary fig. S8, Supplementary Material online). This pattern remained unchanged when only samples homozygous for the minor arrangement were included (fig. 3e; supplementary fig. S9, Supplementary Material online).

When looking at the ratio of nonsense mutations versus nonsynonymous mutations (*P*_nonsense_/*P*_0_), we found a stronger negative correlation with recombination rates in all species (*H. annuus*: Pearson’s r = - 0.2770, p < 2.2×10^−16^; *H. argophyllus*: Pearson’s r = -0.3104, p < 2.2×10^−16^; *H. petiolaris*: Pearson’s r = - 0.2183, p < 2.2×10^−16^; fig. 4a-c) than for π_0_/π_4_. Similar to the results with π_0_/π_4_, however, there was no excess in their proportion within inversions when compared with the genome-wide average for windows with a similar recombination rate (fig. 4d; supplementary fig. S10, Supplementary Material online). One exception was in the “reduced” category in ann15.01 (p=0.0104; fig. 4d), although the increase in the “null” category in the same inversion was not significant. When only minor arrangements were analyzed, the “reduced” category in ann15.01 continued to have greater *P*_nonsense_/*P*_0_ than the genomic average (fig. 4e). Such a significant increase was also found in the “high” category in inversion pet11.01 and in pet14.01, while the *P*_nonsense_/*P*_0_ values did not differ significantly for other categories and inversions (fig. 4e; supplementary fig. S11, Supplementary Material online).

**Fig 4.**
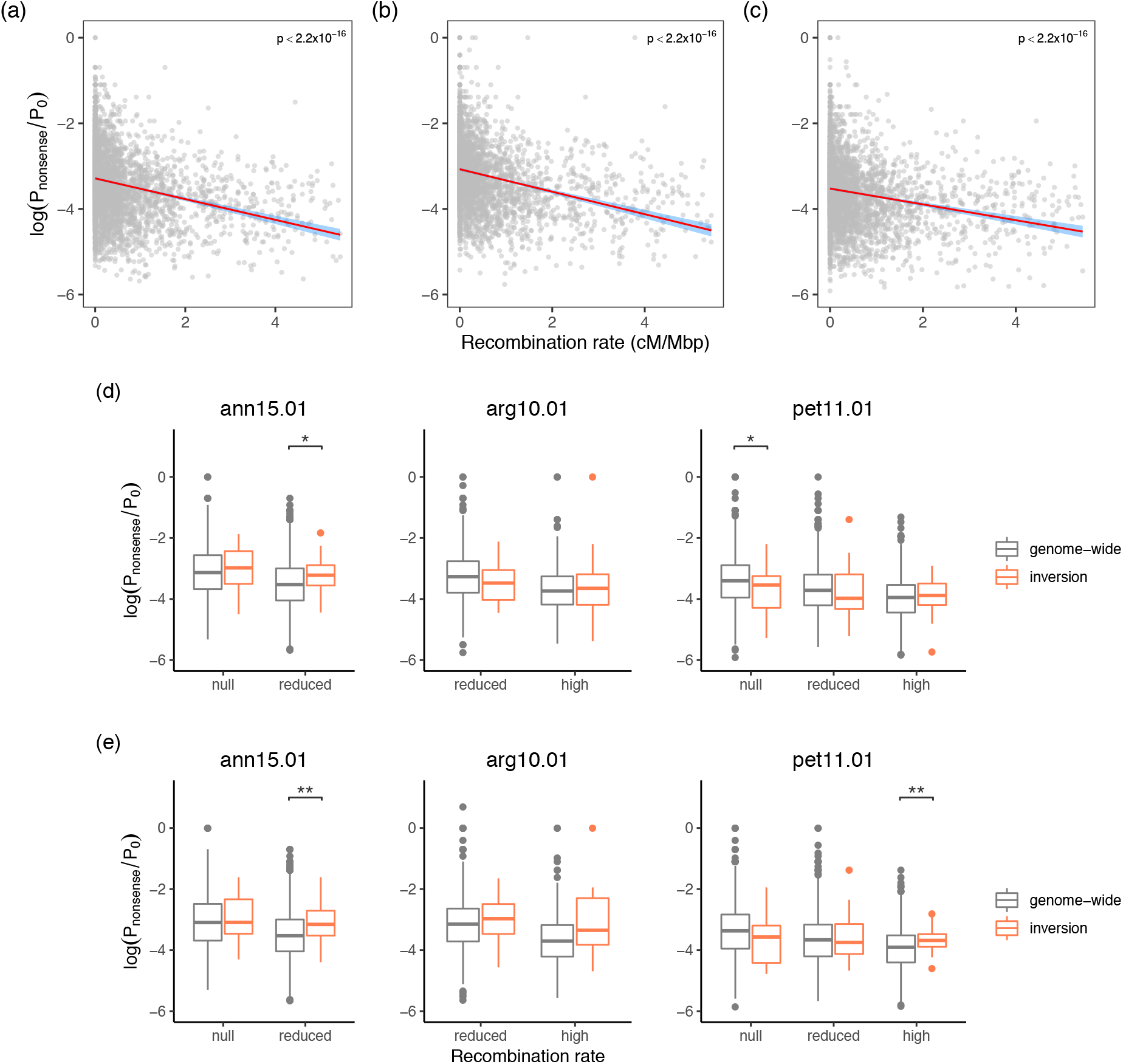
Recombination rate and proportion of nonsense mutations. Genome-wide ratio of nonsense mutations and zero-fold mutations (*P*_nonsense_/*P*_0_) was compared to recombination rate in (a) *Helianthus annuus*, (b) *H. argophyllus* and (c) *H. petiolaris*. The red lines denote the best-fit linear regression line with the 95% confidence intervals shaded in blue. The values within inversions were compared to genome-wide regions in the same recombination rate category using (d) all samples or (e) samples homozygous for the rarer arrangement. Windows of each recombination rate category were compared separately. Asterisks denote significance in independent t-test: *0.05>p>0.01; **0.01>p>0.001. Results of all inversions are presented in supplementary fig. S10 and S11, Supplementary Material online.

To more precisely estimate how genotype frequency of inversions affects the accumulation of deleterious load, we also compared π_0_/π_4_ and *P*_nonsense_/*P*_0_ between samples from populations that contain only one inversion arrangement and samples from populations that are polymorphic. For three inversions pet05.01, pet09.01 and pet11.01, which show very different distribution patterns between subspecies, homozygous samples from populations with both arrangements showed strikingly higher π_0_/π_4_ and *P*_nonsense_/*P*_0_ compared to samples of the same genotype in populations with only one arrangement, when controlled for differences in demography (fig. 5; supplementary fig. S12, Supplementary Material online).

**Fig 5.**
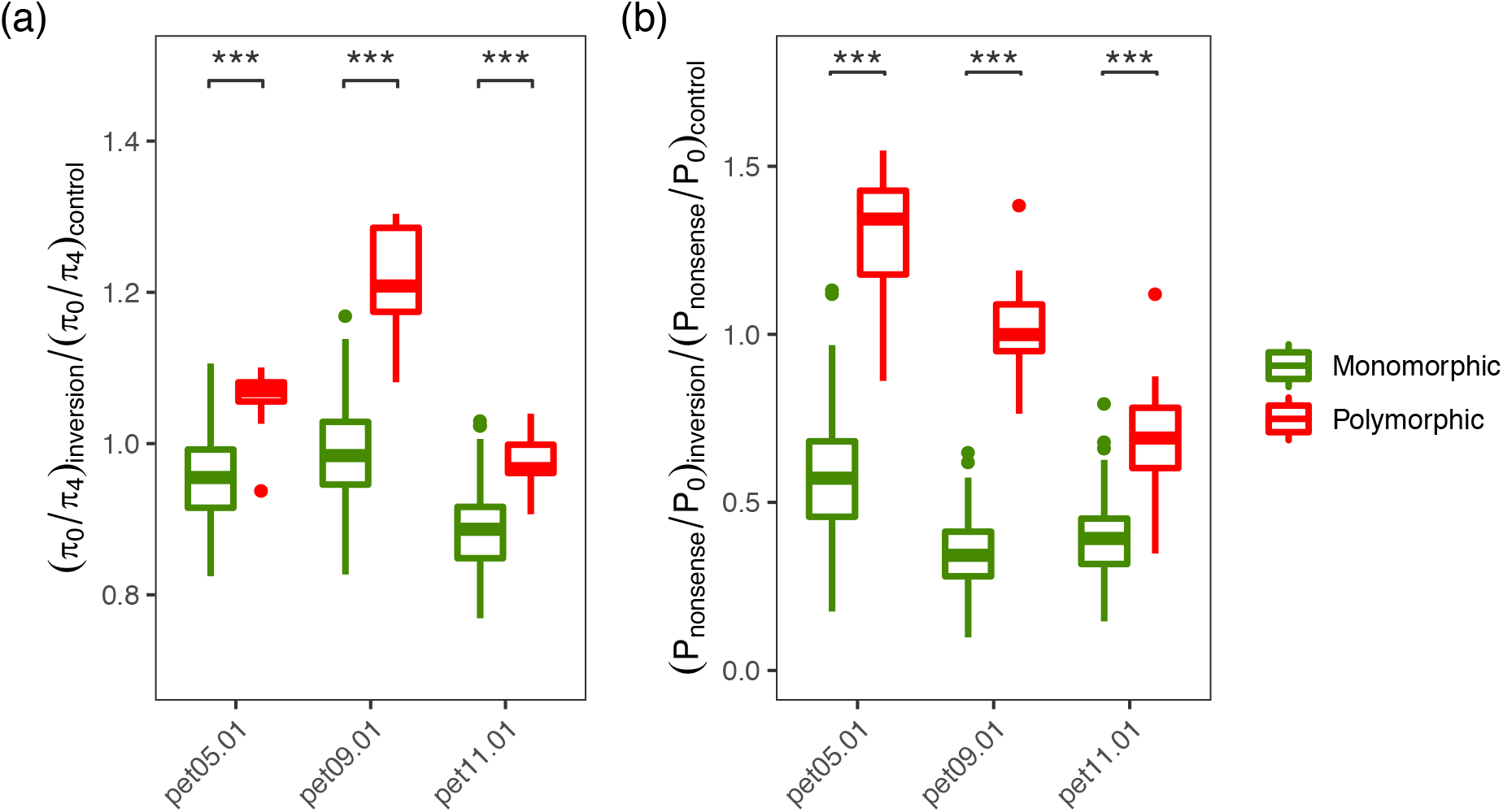
Comparisons of (a) π_0_/π_4_ and (b) *P*_nonsense_/*P*_0_ between monomorphic populations and polymorphic populations for inversions pet05.01, pet09.01, and pet11.01. The statistics were calculated for samples with the same homozygous genotype from populations that were monomorphic and polymorphic for an inversion and normalized to account for differences in demography. Asterisks denote significance in independent t-test: ***p<0.001. Distribution maps of inversion genotypes in populations used in the analyses are presented in supplementary fig. S12, Supplementary Material online.

### Establishment of the inversion polymorphism

Polymorphic inversions can arise from divergent selection or balancing selection (Faria et al. 2019), leading to different predictions regarding fitness of genotypes and, by inference, genotype frequency. In the circumstance of divergent selection, arrangements capturing different sets of locally adapted alleles should be favored in populations with different environmental conditions, and heterokaryotypes are likely to suffer reduced fitness if the alleles are not completely dominant. A balanced polymorphism can establish through favorable epistatic interactions among alleles at different loci in an inversion (Charlesworth and Charlesworth 1973), or through associative overdominance caused by complementation of recessive deleterious alleles by dominant alleles from the alternate arrangement. In the latter case, individuals heterozygous for an inversion should possess higher fitness and occur at higher frequency than expected under Hardy-Weinberg equilibrium. To test whether increased deleterious load in the inversions contribute to the maintenance of their polymorphism (Connallon and Olito 2021; Berdan et al. 2021), we used the extensive records of developmental, morphological and physiological traits for the samples from a previous common garden experiment (Todesco et al. 2020) to search for evidence of overdominance of the inversions. Among the 87 traits across three species, many inversion genotypes did not differ in phenotype. In several cases, however, inversion heterozygotes fell between the homozygous genotypes, such as in ann01.01 for the diameter of the inflorescence disk and in pet05.01 for leaf area, or displayed a similar phenotype to one of the homozygotes, such as in ann13.01 for phyllary width and in pet11.01 for total leaf number. The only cases where heterozygotes have a significantly (p < 0.01) more extreme phenotype were in inversion ann15.01 for phyllary width and phyllary diameter (supplementary fig. S13, Supplementary Material online).

The presence of recessive deleterious alleles unique to one inversion arrangement will result in reduced fitness in homozygotes for the inversion and increase the frequency of heterozygotes. However, in our examination of genotype frequencies, we found no excess of inversion heterozygotes across the three species. Instead, when analyzed species-wide, there was an excess of homozygotes (U-score > 0) for all of the inversions that we examined, and all except one (pet11.01) of the inversions showed significantly higher (p < 0.001) homozygosity than the genomic background level (fig. 6). At the population level, U-scores were highly variable, but almost none were significantly different from Hardy-Weinberg expectations. Two exceptions did occur, where populations show a significant excess of homozygotes (supplementary fig. S14, Supplementary Material online).

**Fig 6.**
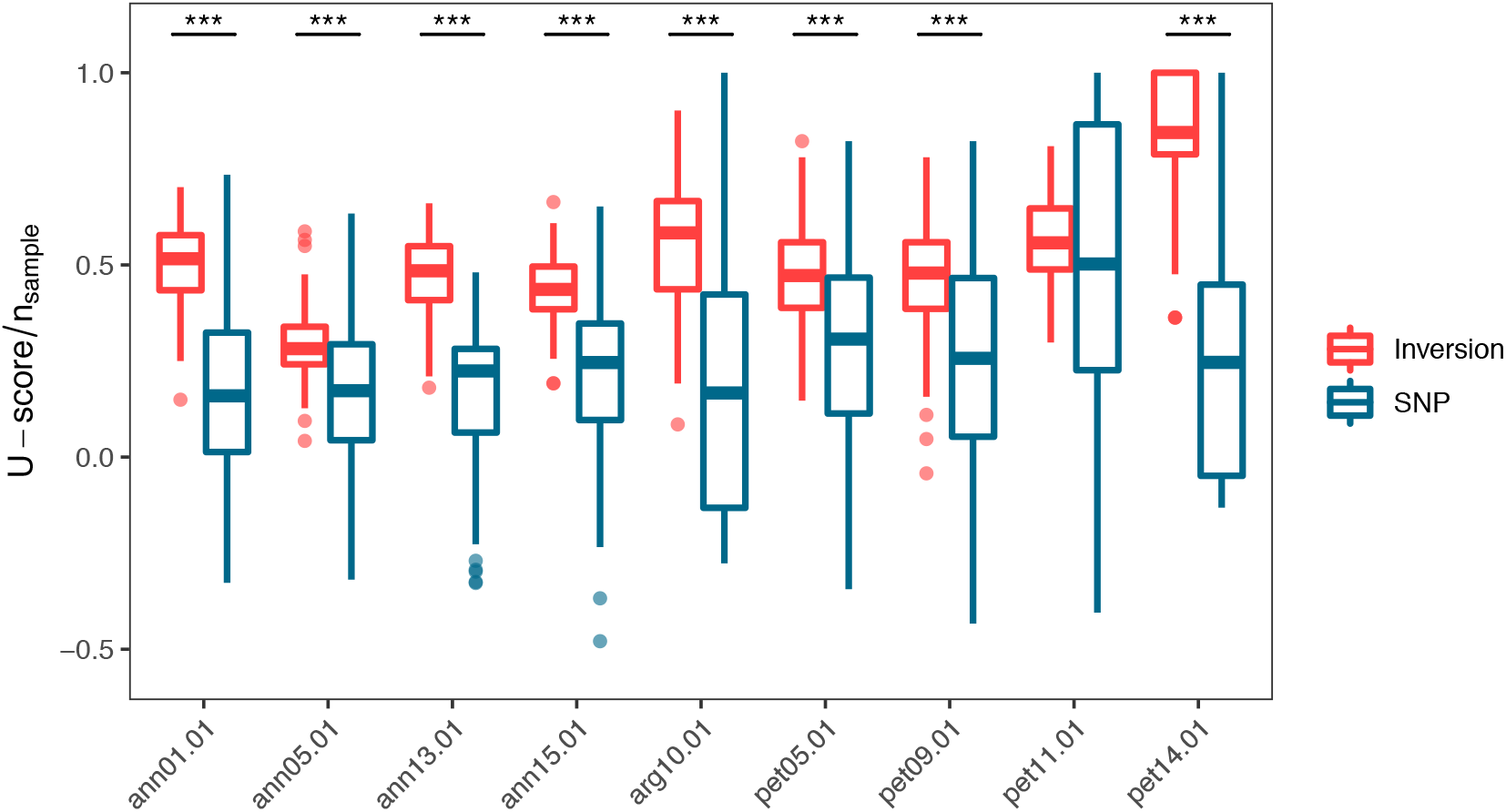
Excess of homozygosity of inversions. U-scores of both inversions and SNPs were calculated across whole species and scaled by number of samples. A positive U-score indicates excess of homozygotes while a negative value indicates overrepresentation of heterozygotes. One individual was randomly chosen from each population and permutated for 100 times to generate sample sets. A hundred SNPs in the same 0.1 allele frequency bin as the inversion and in regions >100kb from all inversions were randomly chosen as a control. Asterisks denote significance in independent t-test: ***p<0.001.

## DISCUSSION

Using three genome assemblies of cultivated sunflower and newly-generated SNP datasets in three wild sunflower species, we analyzed genome-wide patterns of recombination, TE abundance and protein evolution. We found that all of the statistics that we used regarding deleterious mutations, especially *P*_nonsense_/*P*_0_, displayed a significant negative correlation with recombination rate across the genome in all three species. These results are consistent with previous reports of enriched deleterious mutations in genomic regions with reduced levels of recombination (Lu et al. 2006; Renaut and Rieseberg 2015; Lozano et al. 2021). Furthermore, we found a negative correlation of genome-wide π_0_/π_4_ with effective population size in the three species. Reductions in population size and inbreeding will lower effective rates of recombination and reduce the efficacy of natural selection in removing nonadaptive mutations (Charlesworth et al. 1993; Lynch et al. 1993). As a consequence, deleterious mutations should be enriched in taxa with reduced population size, such as asexual species (Hollister et al. 2015), domesticated crops (Lu et al. 2006; Renaut and Rieseberg 2015), and taxa with a history of population bottlenecks (Marsden et al. 2016; Yang et al. 2018; Chen et al. 2020). Our results from the analyses of sunflower genomes are broadly consistent with this earlier work and further support the use of π_0_/π_4_ for measuring the load of deleterious mutations.

Besides protein sequences, we also found an enrichment of LTR-RTs in regions of low recombination in sunflowers. Negative associations between TEs and recombination rates have been commonly observed in eukaryotic genomes (Bartolome et al. 2002; Tian et al. 2009; Gion et al. 2016), although the factors driving these correlations remain poorly understood (Kent et al. 2017). One way in which recombination can affect TE accumulation is through the action of Hill-Robertson interference, assuming that the deleterious effects of TEs mainly result from insertions into genes or regulatory sequences. However, simulations under this model showed that TEs accumulate only in regions of extremely low recombination and when TEs do not excise (Dolgin and Charlesworth 2008). The deleterious effects of TEs can also be caused by ectopic recombination between non-homologous element copies. The ectopic recombination rate is assumed to closely follow the meiotic recombination rate (Kent et al. 2017). Therefore, TEs in actively recombining regions are more likely to generate deleterious chromosomal rearrangements and thus be eliminated by selection. Unequal homologous recombination has been shown to be an important mechanism for the removal of LTR-RTs in sunflowers (Staton et al. 2012), and variation in its incidence across the genome may account for the distribution observed here. Another possible reason for the negative association with recombination is that the insertion of TEs leads to a direct modification of rates of recombination (Kent et al. 2017). However, recombination rate in sunflower species is relatively stable despite difference in TE content (Barb et al. 2014), so the large-scale recombination landscape estimated from genetic maps appears to be largely independent of TEs. Therefore, while variation in effective recombination rates undoubtedly affects the pattern of TE abundance in sunflowers, identification of the specific mechanisms driving TE accumulation in sunflower genomes require further investigation.

While all statistics, including LTR-RT density, π_0_/π_4_ and *P*_nonsense_/*P*_0_, captured the effects of population size and recombination rate variation across the genome, none of the nine locally-adapted inversions that we examined displayed the predicted increase in TE abundance or deleterious amino acid substitutions when compared to other regions of the genome with a similar background recombination rate, except for increases in *P*_nonsense_/*P*_0_ ratio in parts of three inversions (ann15.01, pet11.01 and pet14.01). The finding of an increased *P*_nonsense_/*P*_0_ ratio in parts of the ann15.01, pet11.01 and pet14.01 inversions implies that it is a more sensitive indicator of deleterious load than LTR-RT density or π_0_/π_4_. This observation is consistent with the stronger correlation of the former with recombination rate (fig. 4a-c). On the other hand, these inversions are generally large in size and large inversions are more likely to carry deleterious mutations during their establishment (Santos 1986; Connallon and Olito 2021). In ann15.01 and pet11.01, the increase in *P*_nonsense_/*P*_0_ was found in windows with high or moderate background recombination rate, but not in the “null” categories within the same inversions, possibly suggesting that the impact of recombination suppression may be stronger in those regions. It is noteworthy that for pet14.01 an impact of recombination suppression is only seen in the minor rearrangement, which has the lowest minor allele frequency among inversions examined (table 1). Thus, allele frequency likely plays a critical role in affecting genetic load in these inversions.

Consistent with observations of minimal deleterious load in sunflower inversions, we found little evidence of overdominance of these inversions in the 87 traits examined. There was a hint of heterosis in ann15.01, one of the largest inversions, for phyllary width and diameter. This is consistent with the elevated *P*_nonsense_/*P*_0_ observed in this inversion and with a possible link between inversion lengths and deleterious mutations (Connallon and Olito 2021). For many important traits, the inversions were dominant or incompletely dominant, which does not confer an advantage on heterozygotes. One case of underdominance was found in pet05.01 for internode length, but whether this confers advantage or disadvantage on heterozygotes is unknown. Nevertheless, we only tested a limited set of phenotypes, and we assumed that selection would favor the extrema in a single trait while certain combinations of traits might be the target of natural selection. Common garden experiments in native habitats are needed to investigate the effect of different inversion genotypes on fitness (e.g., Goebl et al. 2020).

Analyses of genotype frequencies of inversions also failed to support the overdominance model. Across the ranges of species, we found an excess of homozygotes for all inversions, which is contrary to expectations for heterosis. A previous examination of inversion genotypes in a dune system in *H. petiolaris* also provided evidence of a heterozygosity deficit for inversions pet05.01, pet09.01 and pet11.01 (Huang et al. 2020). However, population structure may also cause deviations from Hardy-Weinberg equilibrium, so, this result should be viewed as an overall summary of genotypic proportions within species, rather than a strict test of differential mating or survivorship among genotypes within populations. Nevertheless, when compared to SNPs with similar allele frequencies from the rest of the genome, the inversions displayed higher homozygosity, consistent with the presence of selection for inversion homozygotes and/or against heterozygotes, or greater environmental selection. For local populations where gene flow should be unrestricted, most did not have U-scores significantly different from 0, although the sample size for each population was too small for a valid test. In addition, the samples used were collected from seeds on mature plants, and thus reflected post-mating population frequencies rather than those of living plants, so the impact of viability selection could not have been detected (Goebl et al. 2020).

Although the cause for the excess of inversion homozygotes within sunflower species is not fully resolved, multiple lines of evidence have demonstrated that many inversions in sunflowers are under divergent ecological selection. For example, previous studies have associated inversions with numerous environmental variables and locally adapted traits (Huang et al. 2020; Todesco et al. 2020). For instance, in *H. annuus*, inversion ann13.01 was associated with flowering time and climate continentality; in *H. argophyllus*, inversion arg10.01 is enriched on the barrier islands of Texas and was associated with temperature and relative humidity; pet05.01, pet09.01, pet11.01 and pet14.01 were all found to contribute to dune adaptation in *H. petiolaris* (Huang et al. 2020; Todesco et al. 2020). Hybrids between dune and non-dune ecotypes of *H. petiolaris*, which are enriched with different inversion arrangements, showed reduced fitness in either of the local environments in reciprocal transplant experiments (Ostevik et al. 2016), and Goebl et al. (2020) showed that several of these inversions were under divergent natural selection. Such divergent selection could account, in part, for the excess of homozygous genotypes, especially at locations where selection is strong and likely explains the contrary results in sunflower inversions and those in Insecta, many of which establish via associative overdominance (Butlin and Day 1985; Eanes et al. 1992; Sniegowski and Charlesworth 1994; Jay et al. 2021).

Assortative mating may also play a role in maintaining the high homozygosity of inversion arrangement. Such assortative mating can occur via ecogeographic isolation since, as discussed above, sunflower inversions contribute to local adaptation. Conspecific pollen precedence was found between dune and non-dune ecotypes in *H. petiolaris* (Ostevik et al. 2016), although whether the underlying loci are located in inversions is still unknown. However, theoretical studies suggest that this is likely the case since inversions offer a means to establish linkage disequilibrium between loci for local adaption and assortative mating (Trickett and Butlin 1994; Servedio 2009; Huang and Rieseberg 2020).

Our analyses of the deleterious load among populations with differences in genotype composition further confirmed that levels of deleterious load in inversions varies with heterozygote frequencies. In populations where inversions are polymorphic, inversions exhibited higher deleterious load compared to samples from populations where only one arrangement is found (fig. 5). These striking differences between polymorphic and monomorphic populations confirm that recombination suppression in inversion heterozygotes, to some extent, has detrimental consequences in sunflower and that high homozygosity of inversions largely averts such cost. We also expect increased TE abundance in populations with a higher frequency of inversion heterozygotes, but because of the difficulty in precisely identifying TEs within inversions using short reads data, we could not test this hypothesis in the present study.

In addition to high homozygosity of inversions, gene conversion can also mitigate the accumulation of deleterious mutations (Berdan et al. 2021). Our examination of the pattern of *F*_ST_ between karyotypes suggest that while inversions allow independent evolution of chromosomes with different arrangements, there is a visible level of gene flux between arrangements due to double recombination or gene conversion, especially in the largest inversion ann13.01 (supplementary fig. S3, Supplementary Material online). Such flux may further limit deleterious mutation accumulation in the inversions, but precise measurement of gene flux between arrangements requires genetic mapping in specific crosses (Crown et al. 2018; Korunes and Noor 2019).

Taken together, our results suggest that the large inversions we have discovered to be segregating in sunflower populations are generally less prone to deleterious mutation accumulation than expected. This is like mainly due to the high frequency of inversion homozygotes in natural populations, but gene conversion may contribute as well. High homozygosity of inversion arrangements should be common for locally adapted inversions, especially when the environmental differences are correlated with geography and vagility is limited. Theoretical studies of the role of recessive deleterious mutations in inversions normally assume mutation-selection equilibria (Nei et al. 1967; Ohta 1971; Santos 1986; Berdan et al. 2021). Analyses of deleterious mutation accumulation in the common scenario of local adaptation is required for a better understanding of inversion fates.

Recombination increases the rate of adaptive evolution by bringing beneficial mutations together and decoupling them from deleterious mutations, while the suppression of recombination can facilitate adaptive divergence in the face of gene flow by holding advantageous alleles together. Thus, recombination modifiers that simultaneously permit high levels of recombination within populations, while reducing such recombination between ecologically divergent populations should be favored. Our analyses in sunflowers suggest that inversions can offer such versatility; recombination can occur among chromosomes of the same arrangement, but not between arrangements, thereby permitting divergence with gene flow, whilst largely averting the accumulation of deleterious mutations. This contrasts with other genomic features known to suppress meiotic recombination, such as telomeric and heterochromatic regions, which suppress recombination universally. Conversely, several other kinds of structural variants may affect recombination in a similar way as inversions, thus providing an evolutionary advantage. For example, hemizygosity resulting from insertion and deletion polymorphisms also precludes recombination between chromosomes with different haplotypes, but not within those possessing the same haplotype (He and Dooner 2009; Schwander et al. 2014; Lawrence et al. 2017). However, compared to inversions, because deleterious mutations would not be masked in hemizygous genotypes, hemizygosity may follow a different evolutionary trajectory.

## MATERIALS AND METHODS

### Samples and variant calling

Most of the sequences analyzed in the present study were generated by Todesco et al. (2020). In brief, samples were grown from seeds collected from wild populations across the native range of each species. Tissue from young leaves was collected from all individual plants, and genomic DNA was extracted from leaf tissue to prepare paired-end whole-genome shotgun (WGS) Illumina libraries. After treatment with duplex-specific nuclease to reduce the representation of repetitive sequences, the libraries were sequenced on HiSeq2500, HiSeq4000 and HiSeqX instruments to produce paired-end, 150-bp reads. Illumina adapters and poor quality reads were clipped using Trimmomatic v0.36 (Bolger et al. 2014) and reads shorter than 36 bp were discarded. These processes resulted in an average 6.34-fold coverage of gene space (Todesco et al. 2020).

To increase sampling of rare inversion arrangements, we collected and sequenced additional samples from two dune systems in *H. petiolaris*, where a number of inversions are enriched in the dune ecotypes (Todesco et al. 2020). Specifically, in 2008, we visited Great Sand Dunes National Park and Preserve in Colorado, USA and collected seeds from 11 dune and 9 non-dune subpopulations; in 2011, we collected seeds from 8 dune and 10 non-dune subpopulations in Monahans Sandhills State Park in Texas, USA (supplementary table S1, Supplementary Material online). To avoid sequencing recent immigrants in each subpopulation, we selected one maternal family from each subpopulation that had seed sizes within the expected range of that subpopulation (i.e. larger seeds for dune subpopulations and smaller seeds for non-dune subpopulations). We germinated seeds in the lab and collected tissue from young leaves from one individual per maternal family (i.e. one individual per population). High molecular weight DNA was extracted from leaf tissue using a modified CTAB protocol (Murray and Thompson 1980; Zeng et al. 2002). TruSeq gDNA libraries for each individual were prepared and sequenced at Genome Quebec Innovation Center (Montreal, Canada). Each library was split across two lanes of Illumina HiSeq 2000 to produce 100-bp paired-end reads.

All the samples were aligned to a newly released reference genome for *H. annuus* (Ha412Hov2.0), which used Hi-C (Marie-Nelly et al. 2014) for contig and scaffold ordering and was shown to have improved quality (Todesco et al. 2020). The filtered reads were aligned to the new reference genome using NextGenMap v0.5.3 (Sedlazeck et al. 2013) and resulting BAM files were concatenated, sorted, and duplicate-marked using samtools v0.1.19 (Li et al. 2009). Libraries sequenced in multiple lanes were merged with sambamba v0.6.6 (Tarasov et al. 2015), and PCR duplicates were remarked.

Variant calling was performed with the Genome Analysis Tool Kit v 4.1.4.1 (GATK; McKenna et al. 2010). To reduce computational time in the variant calling process, we excluded genomic regions that contain transposable elements (TEs), which represent more than ¾ of the sunflower genome (Badouin et al. 2017), as well as small unplaced chromosome contigs, chloroplast, and mitochondria. For each sample, a GVCF file was produced with the GATK ‘HaplotypeCaller’ with the parameter “--heterozygosity 0.01”. After individual variant calling, all samples from each species were jointly genotyped using GATK’s ‘GenomicsDBImport’ and ‘GenotypeGVCFs’. The step was run over 1 Mbp regions of the genome for parallel computation, and the raw VCF chunks were then gathered by chromosome using ‘GatherVcfs’.

In order to remove low-quality variants, we followed GATK best practices and conducted Variant Quality Score Recalibration (VQSR) to filter the raw VCFs. The 20 samples with the highest sequencing depth in each species were selected to produce a “gold set” using the following parameters: mapping quality > 50.0, missing rate < 10%, -1.0 < strand odds ratio < 1.0, minor allele frequency > 0.25, excess heterozygosity < 10.0, -1.0 < BaseQRankSum < 1.0, depth within one standard deviation from the mean and ExcessHet z-score > -4.5. The raw set of all variants were first filtered to remove sites with extremely heterozygosity (ExcessHet z-score < -4.5) and the gold set was then applied against this filtered set of variants to produce recalibration models for SNPs and indels using ‘VariantRecalibrator’. The 90% tranche for each species was selected based on these recalibration models using ‘ApplyVQSR’. An additional filter was applied for each species to retain only bi-allelic SNPs with minor allele frequency > 0.01 and genotyping rate > 50%.

### Selection of inversions

For this study, we selected a subset of haploblocks from Todesco et al. (2020) for which there was solid evidence of an underlying inversion and found to be important in ecological adaptation. In *H. annuus*, we selected inversions ann01.01 and ann05.01, the structure of which are well-characterized by comparative genomic analysis, plus inversions ann13.01 and ann15.01, which are the largest inversions found in sunflowers (Todesco et al. 2020). In *H. argophyllus*, inversion arg10.01 was chosen because it was shown to be associated with the formation of the early flowering ecotype on the barrier islands of Texas (Moyers 2015; Todesco et al. 2020). In *H. petiolaris*, we selected 4 inversions (pet05.01, pet09.01, pet11.01, pet14.01), which were found to contribute to dune adaptation in the species (Huang et al. 2020; Todesco et al. 2020). Among these inversions, pet05.01, pet09.01 and pet11.01 were used for the comparison of genetic load between homozygous and polymorphic populations, because these inversions show very different distribution patterns between subspecies.

### Estimation of recombination rate and localization of centromeres

We remapped the markers of an integrated genetic map for cultivated sunflower (Badouin et al. 2017) to our current reference genome Ha412Hov2.0 using BWA MEM (Li 2013) with default settings. A cubic smoothing spline was fit to the remapped marker positions on each chromosome and outliers with residual > 1 were removed. We manually inverted the first 10 Mbp of chromosome 12 in the genetic map that was likely due to assembly error, and we fit monotonic increasing genetic coordinates at a resolution of 1 Mbp using a smoothing spline model. Recombination rate was calculated for each 500-kbp non-overlapping window based on the physical and genetic distances on the transposed genetic map.

To determine the location of sunflower centromeres, we made use of the sequences previously found to be targeted by sunflower centromere-specific histone H3 (Nagaki et al. 2015). Four main sequences from Nagaki et al. (2015) were downloaded from the Nucleotide database of National Center for Biotechnology Information (NCBI) with the following accession: LC075744, LC075745, LC075746, and LC075747. These sequences were then queried against the Ha412Hov2.0 genome using BLASTN (https://blast.ncbi.nlm.nih.gov/) with an E-value of 1×10^−5^.0

### Linkage disequilibrium and sequence divergence

To estimate the extent of recombination suppression caused by the inversions, we calculated linkage disequilibrium (LD) between SNPs across the chromosomes where the inversions reside. We removed SNPs with minor allele frequencies < 10%, thinned the variants to one per 5 kbp and calculated pairwise LD (R^2^) values between SNPs using PLINK v1.9 (Chang et al. 2015; Purcell et al. 2007). For each inversion, we calculated LD for all samples from populations that are polymorphic for the inversion arrangements with a minor allele frequency > 0.1 and for samples from populations that contain only the homozygous genotype for the major arrangement (Todesco et al. 2020). Values of SNPs were grouped into 500-kbp windows and the average R^2^ value was calculated for each window pair.

To provide an additional assessment of the effect of inversions in suppressing recombination between arrangements, we calculated Weir and Cockerham’s *F*_ST_ (Weir 1996) in 1-Mbp sliding windows with a step size of 200 kbp and 100-kbp sliding windows with a step size of 20 kbp, respectively, between samples homozygous for each arrangement for each inversion using VCFtools (Danecek et al. 2011) with the SNPs generated on the new reference genome.

### Transposable element abundance

We conducted de-novo TE annotation for the three reference genomes of cultivated sunflowers using the program EDTA (Ou et al. 2016). The program was run with the default settings “--sensitive 0 -- evaluate 0” and parameter “--anno 1” to perform whole-genome TE annotation on the 17 chromosomes of each genome assembly, and known coding sequences of sunflower from the gene annotation of the Ha412HOv2.0 genome (https://sunflowergenome.org/) were provided for TE filtration. We extracted LTR-RTs from the EDTA-generated annotations for downstream analyses because these elements are known to be transcriptionally active in sunflower species (Cavallini et al. 2010; Kawakami et al. 2011; Renaut et al. 2014).

We summarized the density of LTR-RTs by calculating the proportion of structurally intact and fragmented elements (including solo-LTRs and truncated LTR-RTs) in sliding windows of 500 kbp across the genome. The TE densities were then compared with the recombination rates calculated on the Ha412HOv2.0 genome. Because recombination rates are highly heterogeneous across the genome, the accumulation of TEs within an inversion likely depends in part on the local recombination environment. To control for the effect of genomic context on TE density, we divided the windows into three different categories based on the recombination rate: (1) high recombination rate for windows with recombination rate > 2 cM/Mbp, (2) reduced recombination rate regions with recombination rate between 0.01 and 2 cM/Mbp, and null recombination regions with recombination rate < 0.01 cM/Mbp (3) (Bartolome et al. 2002). For each inversion, windows of each category were compared to those of the same category from across the genome. TE proportions were logit-transformed and compared using Welch two sample t-tests. Locations of ann01.01 and ann05.01 on each reference genome were extracted based on previous alignments of the genome assemblies (Todesco et al. 2020).

### Protein evolution

In addition to TEs, we examined if inversions show a greater accumulation of deleterious mutations in protein coding genes relative to background levels. To detect signatures of relaxed negative selection, we explored the pattern of protein evolution using the coding sequences of all protein-coding genes from a recently generated gene annotation for the Ha412HOv2.0 reference genome. In brief, the annotation was produced using the program EuGene (Foissac et al. 2008) and the predicted gene sequences were filtered by removing genes that were aligned to a known TE with > 90% identity and over 80% of its length (https://sunflowergenome.org/). The remaining gene set contains 58,234 genes, 53,800 of which are protein-coding genes. We further removed the coding sequences that do not begin with the regular “ATG” start codons or end with “TAG/TGA/TAA” stop codons or those whose lengths are not a multiple of three, leaving a total of 53,302 genes which were used for all downstream analyses.

We first calculated the ratio of zero-fold to four-fold diversity π_0_/π_4_ (Marsden et al. 2016), which estimates the proportion of amino acid substitutions that are not removed by selection. The zero and four-fold degenerate sites were identified by iterating across all four possible bases at each site along the coding sequences and comparing the resulting amino acids. Sites where the four different bases resulted in four different amino acids were classified as zero-fold degenerate and those with no changes in amino acids were defined as four-fold degenerate. The diversity at zero-fold or four-fold degenerate sites was calculated as:

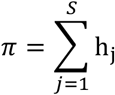

where *S* is the number of segregating sites and *h*_j_ is the heterozygosity for the *j*^th^ segregating site defined as:

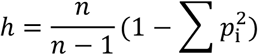

where *n* is the number of haplotype sequences in the sample and *p*_i_ is the sample frequency at the *i*^th^ allele (Tajima 1989; Hahn 2019). The ratio of π_0_ and π_4_ was then calculated in sliding windows of 500 kbp using samples from all populations in each of the three species, or samples homozygous for the minor arrangement. The ratios were then compared to recombination rates and the values for the inversions were compared to genome-wide measures in a similar way to that in the above analysis of TE density, except that the ratios were log-transformed and recombination rate categories with less than 10 windows in an inversion were removed due to low statistical power. We also calculated π_0_/π_4_ for individuals from populations that were polymorphic for an inversion, as well as from those that were monomorphic, to more precisely estimate inversion effects on the accumulation of deleterious load. Specifically, we chose samples from populations that were homozygous for one arrangement and those of the same homozygous genotype from populations polymorphic with a minor allele frequency > 0.2 and compared π_0_/π_4_ between these samples. To control for differences in demography, a region of the same size as the inversions on chromosome 4, where no inversions have been identified in sunflowers, was randomly chosen and the statistics were calculated with the same sets of samples. In addition, we randomly selected 20 samples from each species and calculated π_0_/π_4_ across the whole genome to estimate the deleterious load among species.

We further examined stop codon mutations using the same protein-coding gene set. We annotated the VCFs using the program snpEff v 5.0c (Cingolani et al. 2012) and extracted “nonsense” mutations (alternate stop codon) for each species. The deleterious load was represented as the ratio of the number of nonsense mutations *P*_nonsense_ and the number of mutations at zero-fold sites *P*_0_, which was used as an approximation of nonsynonymous mutations. We calculated this statistic in sliding windows of 500 kbp and compared the inversions to genome-wide measures in the same way as for π_0_/π_4_. We also compared *P*_nonsense_/*P*_0_ between monomorphic and polymorphic populations in the same way as for π_0_/π_4_.

### Overdominance of inversions

We used the phenotypic data of the samples collected in a previous common garden experiment (Todesco et al. 2020) to test whether inversions show overdominance in the wild population. In total, we tested 86, 30 and 69 traits for *H. annuus, H. argophyllus* and *H. petiolaris*, respectively (supplementary fig. S13, Supplementary Material online). The trait measurements were normalized and the averages of inversion heterozygotes were compared to those of each homozygous genotype. Significance was assessed using two-sample t-test if values of heterozygotes were greater or lower than both homozygous groups.

### Genotype frequency

We used the genotype information of the samples from Todesco et al. (2020) to calculate allele frequency and genotype frequency of the inversions and tested for Hardy-Weinberg equilibrium. We calculated U-score for each inversion using the full-enumeration method of “hwx.test” in the R package “HWxtest” v1.1.9 (Engels 2009). A positive U-score indicates excess of homozygotes while a negative value indicates overrepresentation of heterozygotes. We calculated this score using all samples across the species range as well as for each population. For the species-wide calculation, we randomly selected 1 individual from each population for each inversion and repeated the process for 100 times to estimate U-score, and we randomly chose 100 bi-allelic SNPs in the same 0.1 allele frequency bin as the inversion and in regions >100kb away from all inversions to account for possible population structure.

## Acknowledgments

We thank Jean-Sébastien Légaré for help in SNP calling, Shujun Ou and Zhigui Bao for help with EDTA, Jeffrey Ross-Ibarra for helpful discussion on inversions, as well as Darren Irwin, Amy Angert, Keith Adams, Sally Aitken and Marc Johnson for their careful and critical reading of an earlier version of the manuscript. This work was supported by a China Scholarship Council scholarship (no. 201506380099 to K.H.) and an NSERC Discovery grant (no. 327475 to L.H.R.).

## Author Contributions

K.H. and L.H.R. conceived the study; M.T. and N.B. contributed sequence data; K.O. collected and produced sequence data for new samples of *Helianthus petiolaris*; K.H. performed SNP calling, TE annotation and analyzed the data; C.E. mapped the centromeres; G.L.O. helped with analyses; K.H. and L.H.R. wrote the paper; and all authors approved the final manuscript.

## Data Availability

The Ha412HOv2.0, XRQv2, PSC8 genome assemblies are available at https://sunflowergenome.org/ and https://heliagene.org/. The newly generated SNP datasets are available at http://www.helianthome.org/. The trait data used in the overdominance analysis are available at https://easygwas.ethz.ch/gwas/myhistory/public/20/, https://easygwas.ethz.ch/gwas/myhistory/public/21/, https://easygwas.ethz.ch/gwas/myhistory/public/22/, https://easygwas.ethz.ch/gwas/myhistory/public/23/.

Sequence data and codes associated with this paper are available upon request and will be set to SRA and GitHub before publication, respectively.

