## supplementary fig. S1, Supplementary Material online for "High homozygosity of inversions in sunflower species largely averts accumulation of deleterious mutations"

**Table S1.** Sample information for the newly-sequenced individuals in this study.

| Population | Sample ID | Ecotype | Coordinate |
|---|---|---|---|
| GSD | GSD983 | non-dune | 37.663 N 105.625 W |
| | GSD1009 | dune | 37.774 N 105.576 W |
| | GSD1039 | dune | 37.787 N 105.569 W |
| | GSD1085 | non-dune | 37.774 N 105.598 W |
| | GSD1139 | dune | 37.818 N 105.597 W |
| | GSD1169 | non-dune | 37.836 N 105.603 W |
| | GSD1184 | dune | 37.773 N 105.553 W |
| | GSD1258 | dune | 37.786 N 105.531 W |
| | GSD1276 | dune | 37.764 N 105.524 W |
| | GSD1329 | dune | 37.745 N 105.546 W |
| | GSD1375 | non-dune | 37.716 N 105.533 W |
| | GSD1503 | non-dune | 37.765 N 105.615 W |
| | GSD1573 | dune | 37.761 N 105.57 W |
| | GSD1707 | dune | 37.803 N 105.524 W |
| | GSD1732 | dune | 37.807 N 105.554 W |
| | GSD1795 | non-dune | 37.813 N 105.515 W |
| | GSD2008 | non-dune | 37.757 N 105.507 W |
| | GSD2031 | non-dune | 37.758 N 105.509 W |
| | GSD2067 | dune | 37.758 N 105.515 W |
| | GSD2252 | non-dune | 37.672 N 105.592 W |
| MON | MON014 | dune | 31.63151 N 102.80999 W |
| | MON069 | dune | 31.63679 N 102.81361 W |
| | MON106 | dune | 31.64052 N 102.81797 W |
| | MON151 | dune | 31.64075 N 102.81345 W |
| | MON220 | dune | 31.62984 N 102.81567 W |
| | MON263 | non-dune | 31.61642 N 102.81197 W |
| | MON308 | non-dune | 32.3204 N 103.82318 W |
| | MON365 | non-dune | 32.21017 N 103.58173 W |
| | MON401 | non-dune | 32.17832 N 103.38889 W |
| | MON459 | non-dune | 32.0889 N 103.17894 W |
| | MON520 | dune | 32.03659 N 103.15085 W |
| | MON564 | dune | 32.03527 N 103.15001 W |
| | MON606 | non-dune | 31.99352 N 103.13029 W |
| | MON658 | non-dune | 31.82584 N 103.07813 W |
| | MON713 | non-dune | 31.64043 N 102.98539 W |
| | MON768 | non-dune | 31.58296 N 102.87074 W |
| | MON818 | non-dune | 31.61138 N 102.83128 W |
| | MON862 | dune | 31.62488 N 102.81174 W |

GSD: Great Sand Dunes National Park and Preserve; MON: Monahans Sandhills State Park.

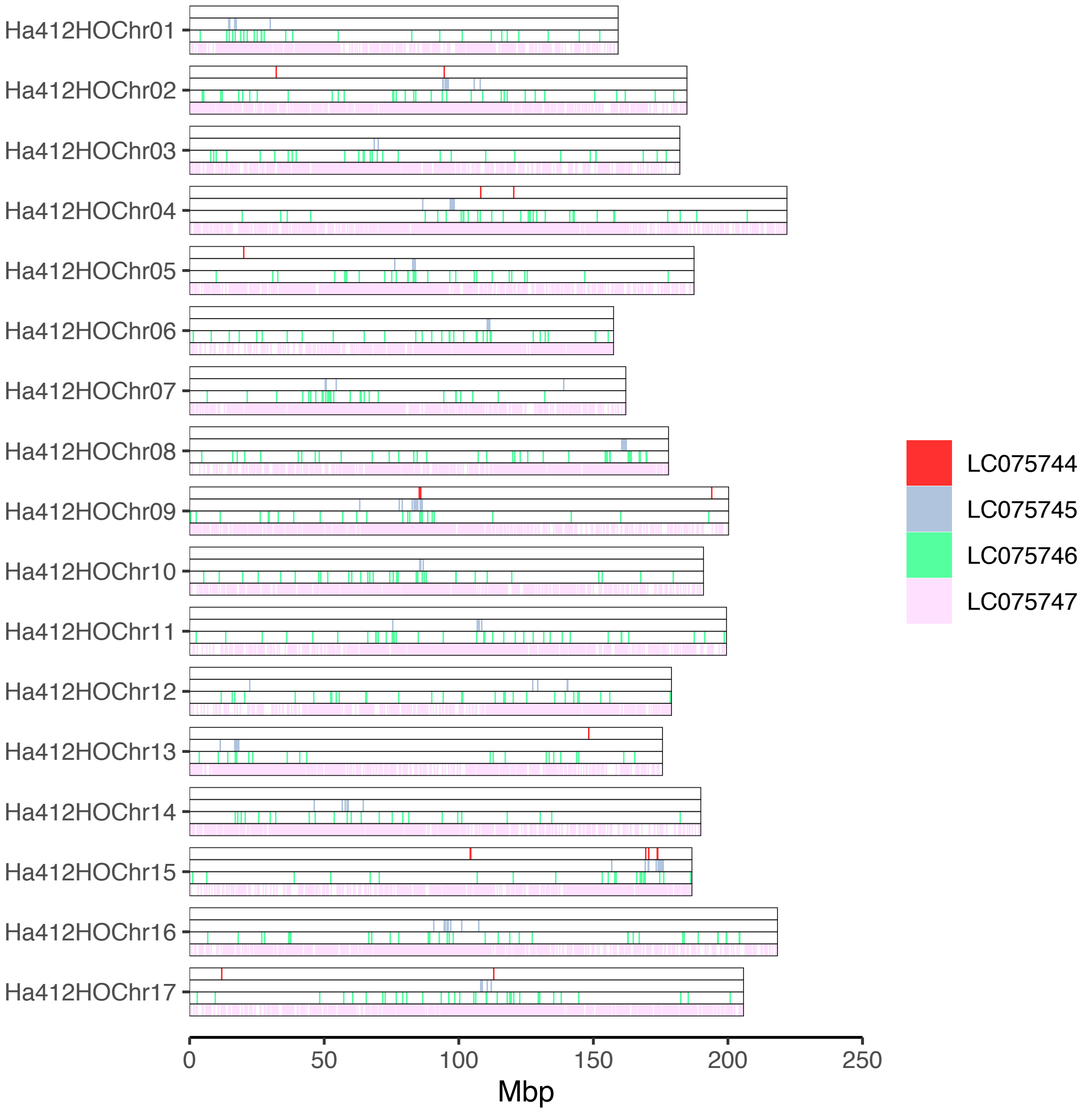


**Figure S1.** Distribution of the sequences previously found to be targeted by sunflower centromere-specific histone H3 across the 17 chromosomes of Ha412HOv2.0.

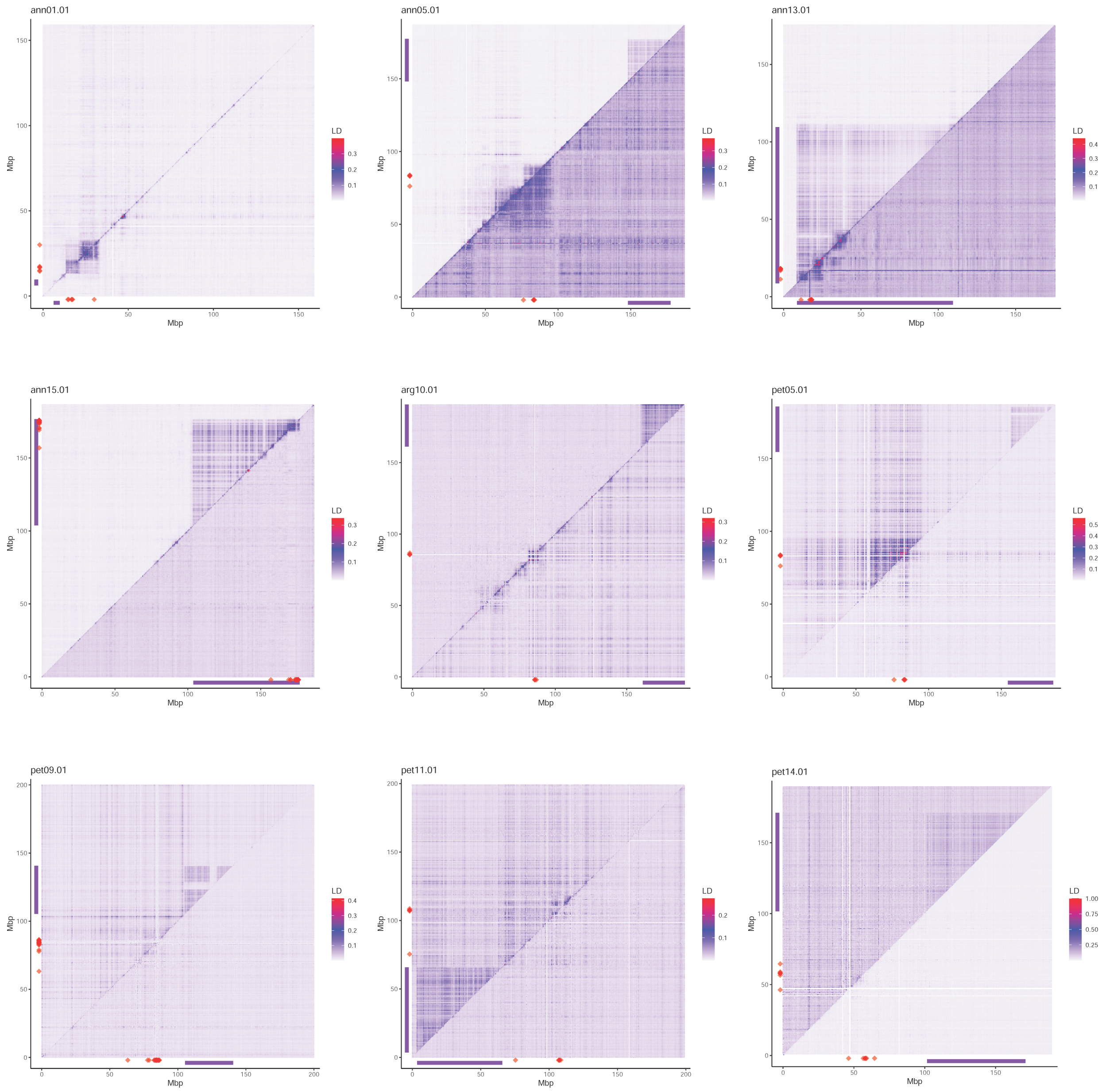


**Figure S2.** LD plots for all inversions. Upper triangle was based on all individuals from populations polymorphic for the inversion and lower triangle was based on individuals from populations homozygous for the more common orientation. Pairwise $R^2$ between SNPs were summarized and the average $R^2$ values were presented in 500-kbp windows. Purple bars represent the location of the inversion. Positions of the centromere-specific sequence LC075745 are indicated by red diamonds. Names for the inversions are consistent with those in Todesco et al. (2020).

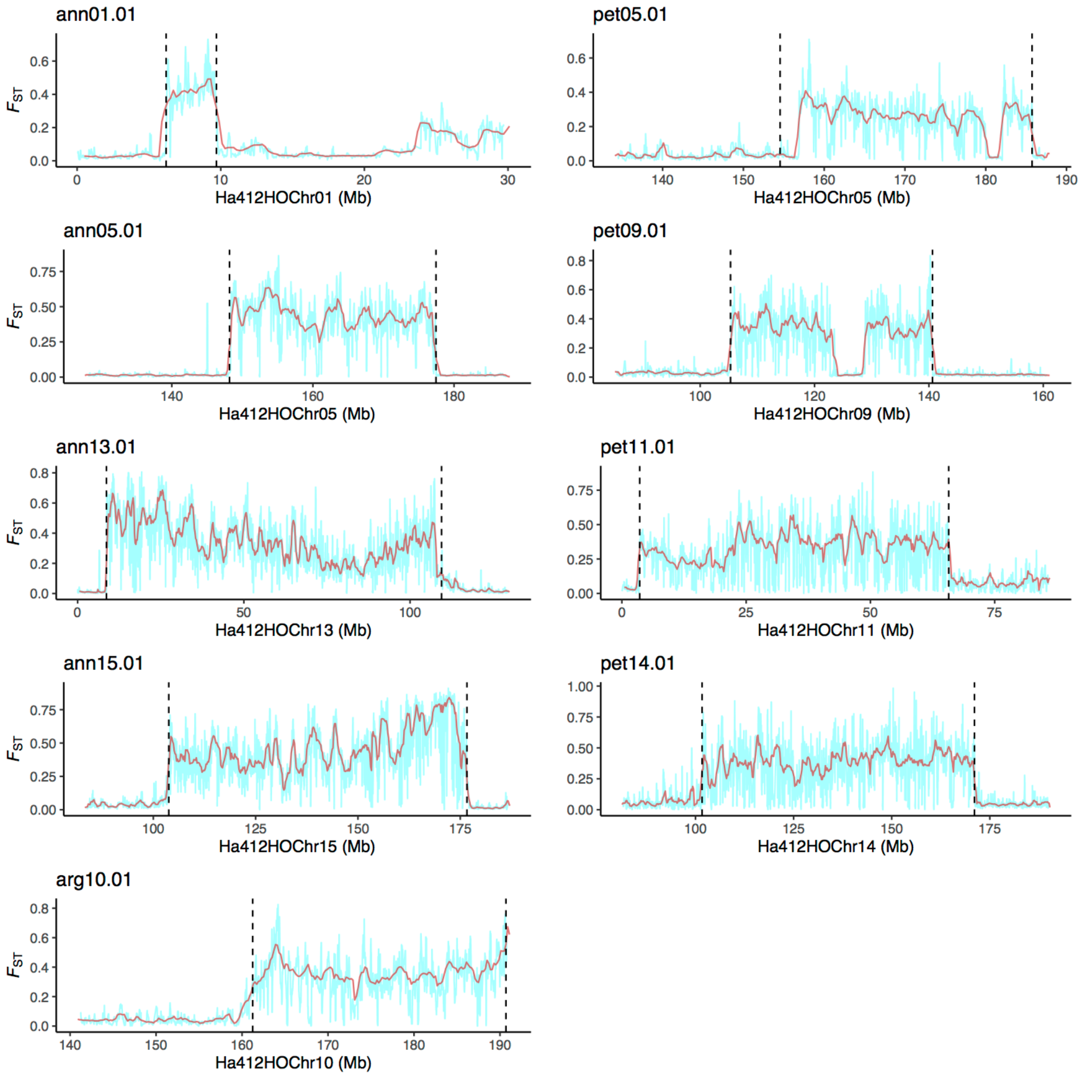


**Fig. S3.** Divergence ($F_{ST}$) between inversion orientations. Mean divergence is calculated over 1-Mbp sliding windows with a step size of 200-kbp (brown lines) and 100-kbp sliding windows with a step size of 20 kbp (cyan lines). Vertical dashed lines indicate the putative boundaries of each inversion.

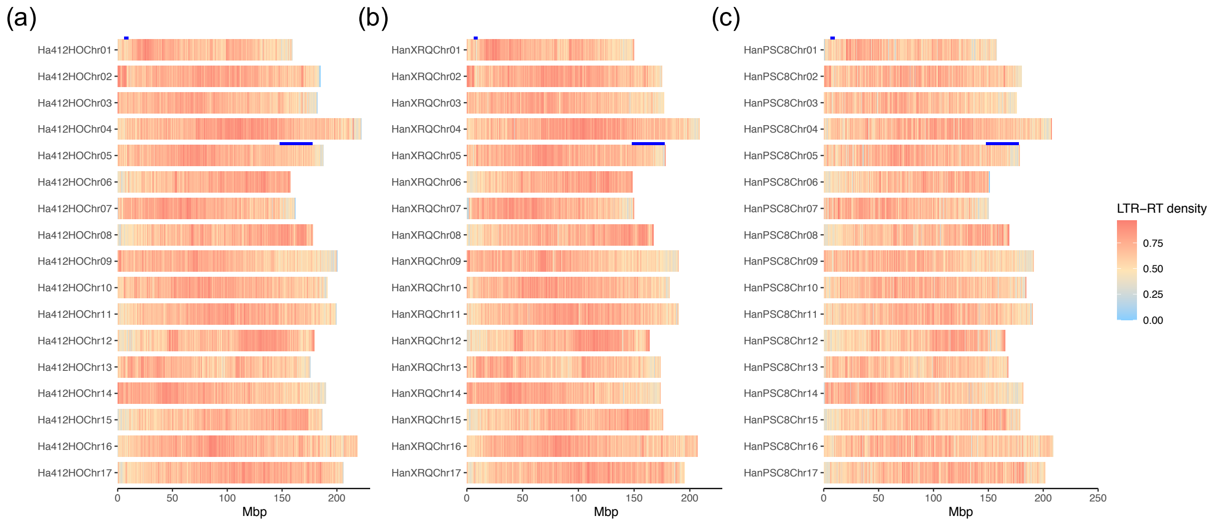


**Figure S4.** Genomic distributions of long terminal repeat retrotransposons (LTR-RTs) in the three genome assemblies of sunflower. (a) Ha412HOv2.0, (b) XRQv2, (c) PSC8. Density of LTR-RTs was calculated in 500 kbp bins per chromosome. Blue bars indicate the locations of inversions ann01.01 and ann05.01.

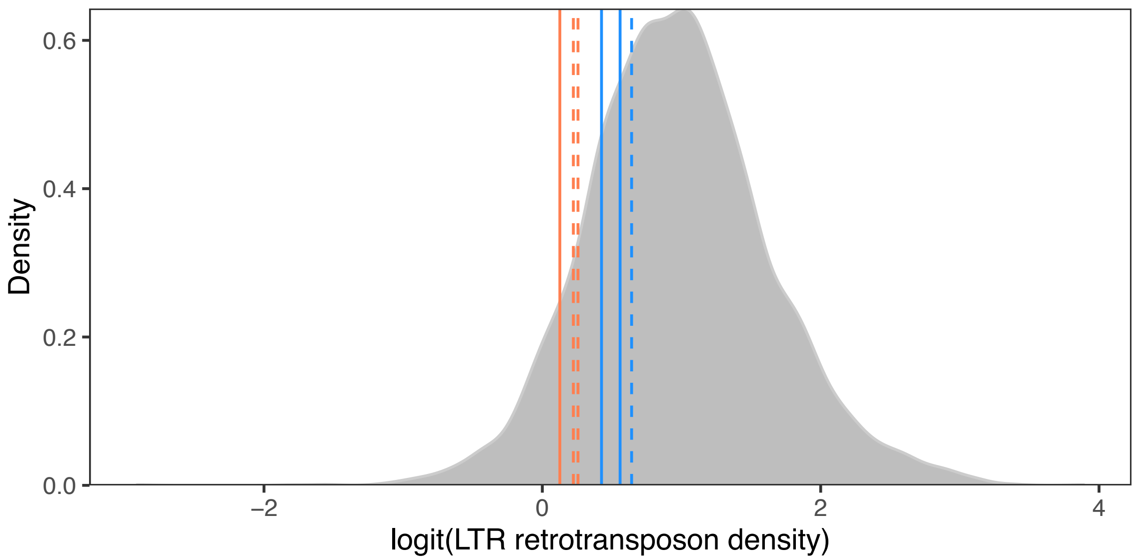


**Figure S5.** Long terminal repeat (LTR) retrotransposon density, computed on 500kb windows genome-wide in Ha412HOv2 reference genome (grey) and in the arrangements of inversion ann01.01 (coral) and ann05.01 (blue) in Ha412HOv2, XRQv2 and PSC8 reference genomes. Solid and dashed lines represent alternative arrangements of the inversions.

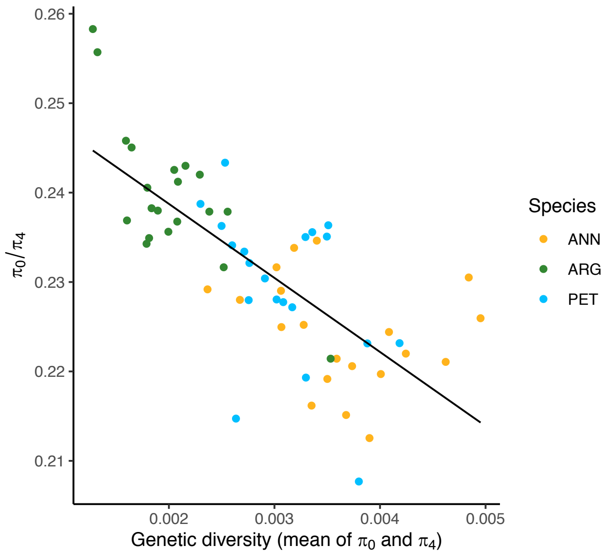


**Figure S6.** Genetic load ($\pi_0/\pi_4$) versus genetic diversity (mean of $\pi_0$ and $\pi_4$) across three sunflower species. The statistics were estimated with 20 individuals randomly chosen from each species of *Helianthus annuus* (ANN), *H. argophyllus* (ARG) and *H. petiolaris* (PET), respectively. The black line denotes the best-fit linear regression line. Both $\pi_0$ and $\pi_4$ were scaled by dividing by the standard deviation of each before the significance of the negative correlation between the variables represented by both axes was tested.

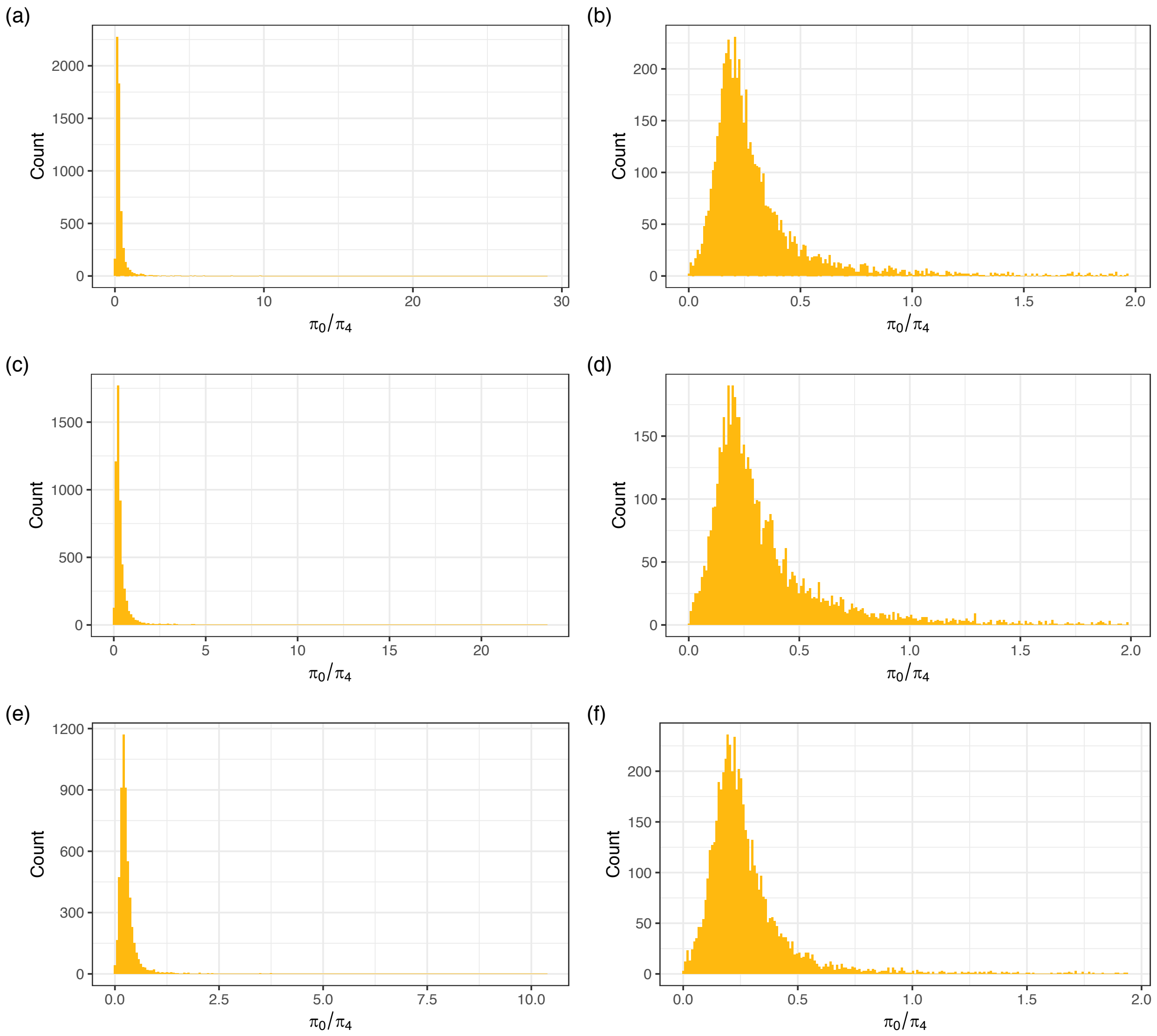


**Figure S7.** Distribution of window-based $\pi_0/\pi_4$ in each species. (a-b) *Helianthus annuus*, (c-d) *H. argophyllus*, (e-f) *H. petiolaris*. The ratio of $\pi_0$ and $\pi_4$ was calculated in sliding windows of 500 kbp. (a), (b) and (c) are distributions for all windows across the genomes**;** (b), (d) and (f) are distributions for windows with values < 2.

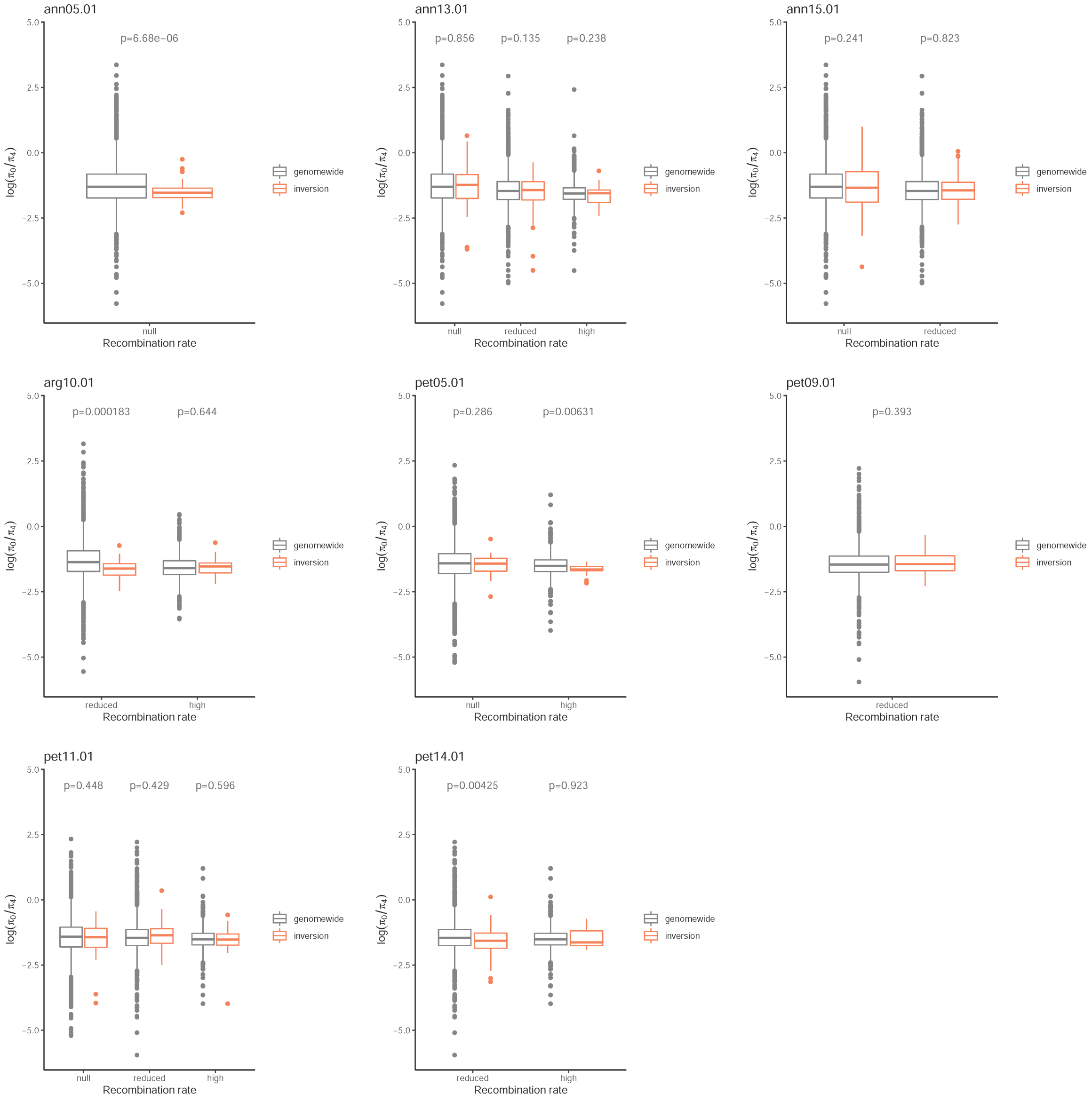


**Figure S8.** Comparison of $\pi_0/\pi_4$ within inversions to genome-wide regions in the same recombination rate category using all samples. Windows of each recombination rate category were compared separately. P-values are Welch two sample t-test on log-transformed $\pi_0/\pi_4$ values.

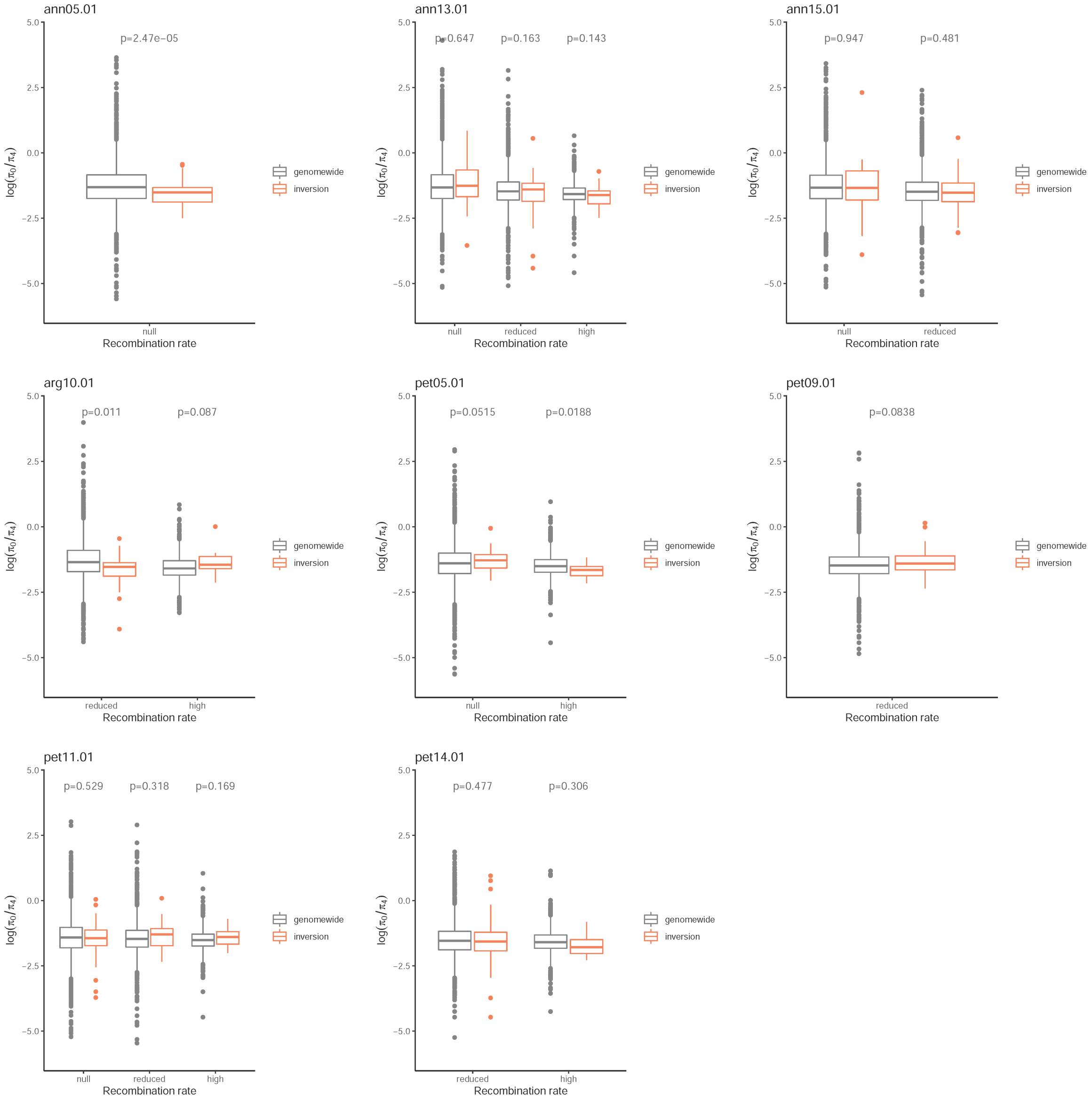


**Figure S9.** Comparison of $\pi_0/\pi_4$ within inversions to genome-wide regions in the same recombination rate category using samples homozygous for the rarer arrangement. Windows of each recombination rate category were compared separately. P-values are Welch two sample t-test on log-transformed $\pi_0/\pi_4$ values.

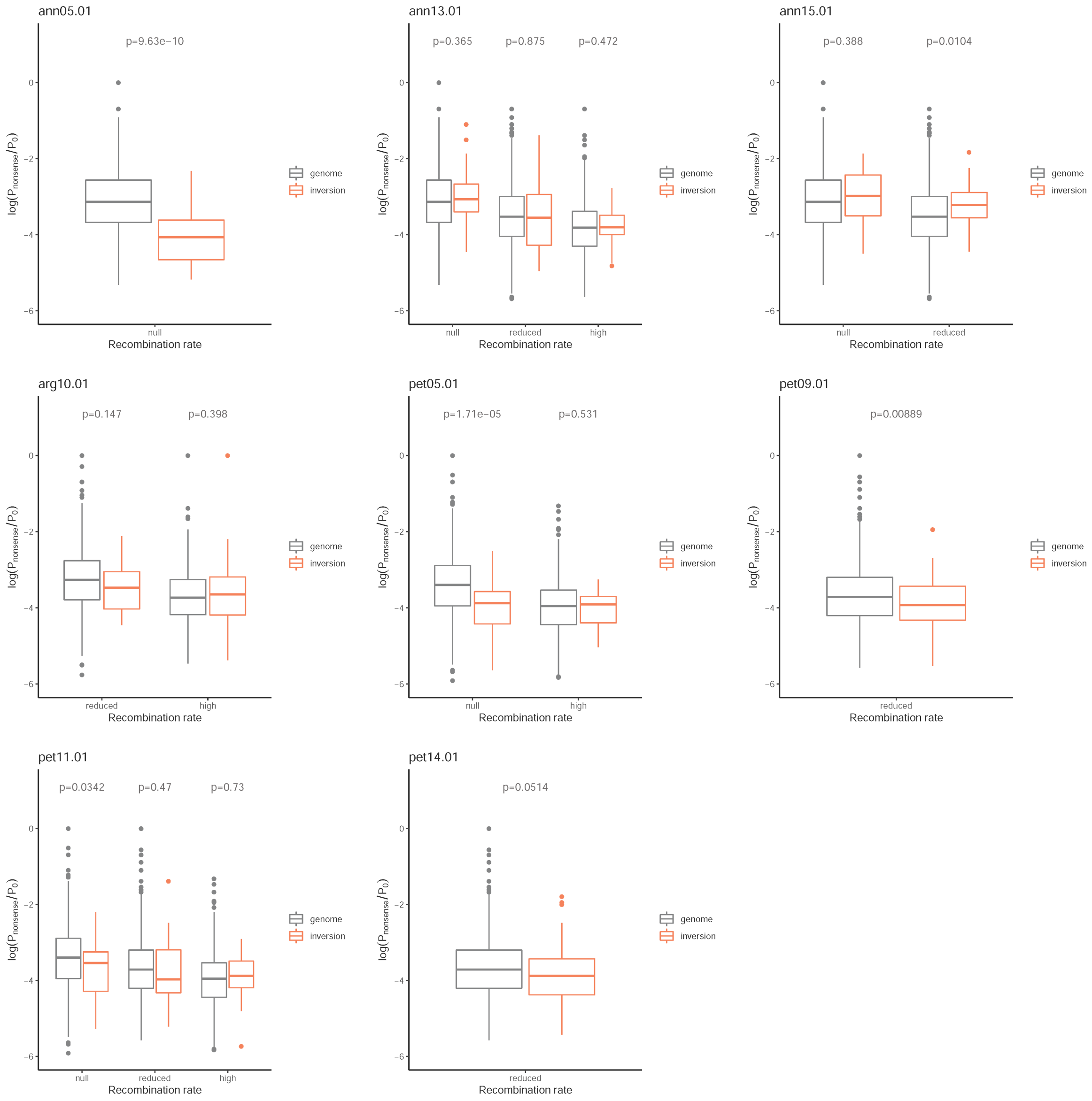


**Figure S10.** Comparison of $P_{\text{nonsense}}/P_0$ within inversions to genome-wide regions in the same recombination rate category using all samples. Windows of each recombination rate category were compared separately. P-values are Welch two sample t-test on log-transformed $P_{\text{nonsense}}/P_0$ values.

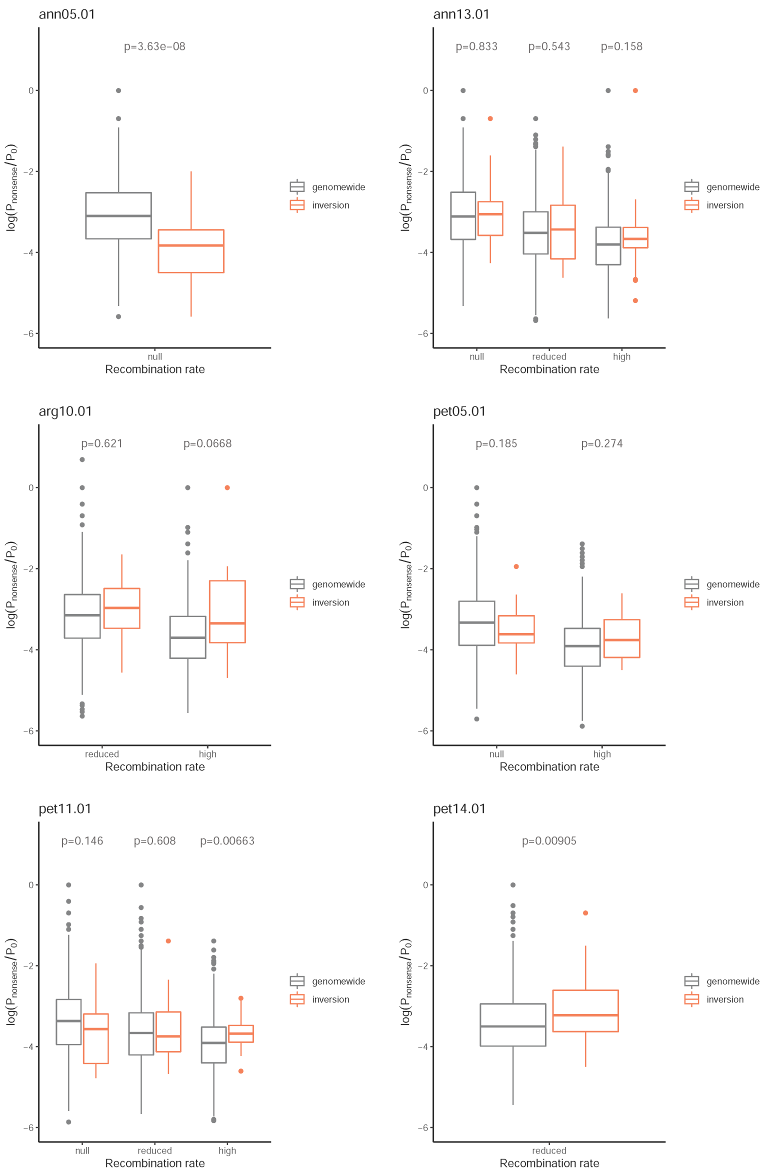


**Figure S11.** Comparison of $P_{nonsense}/P_0$ within inversions to genome-wide regions in the same recombination rate category using samples homozygous for the rarer arrangement. Windows of each recombination rate category were compared separately. P-values are Welch two sample t-test on log-transformed $P_{nonsense}/P_0$ values.

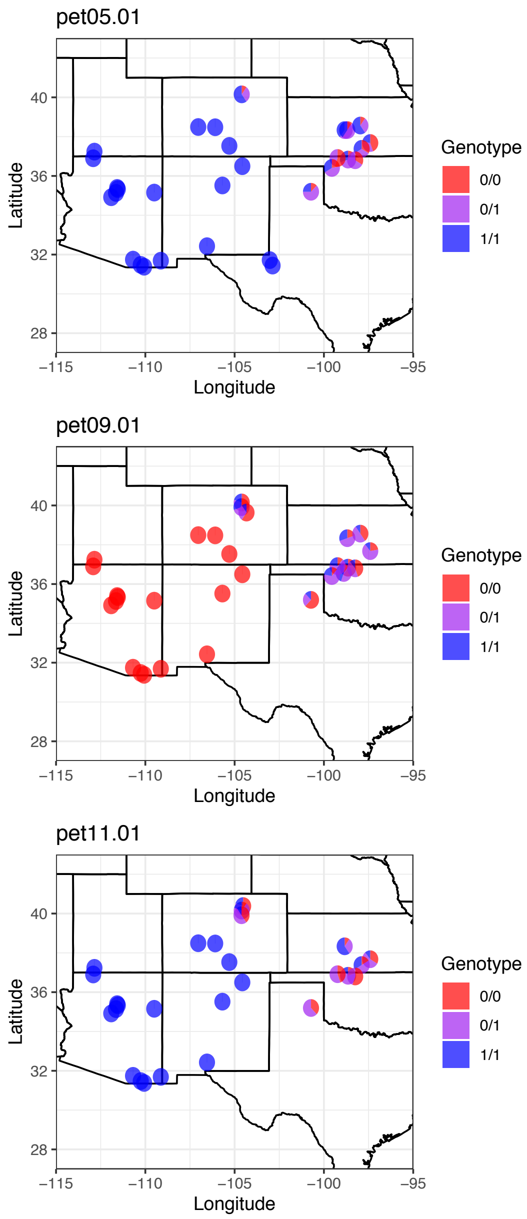


**Figure S12.** Distribution maps of inversion genotypes in populations used in the comparison of deleterious mutations between monomorphic populations and polymorphic populations for inversion pet05.01, pet09.01, and pet11.01. Monomorphic populations contain one arrangement for the inversion (genotype 0/0 or 1/1), and polymorphic populations contain multiple genotypes for the inversions.

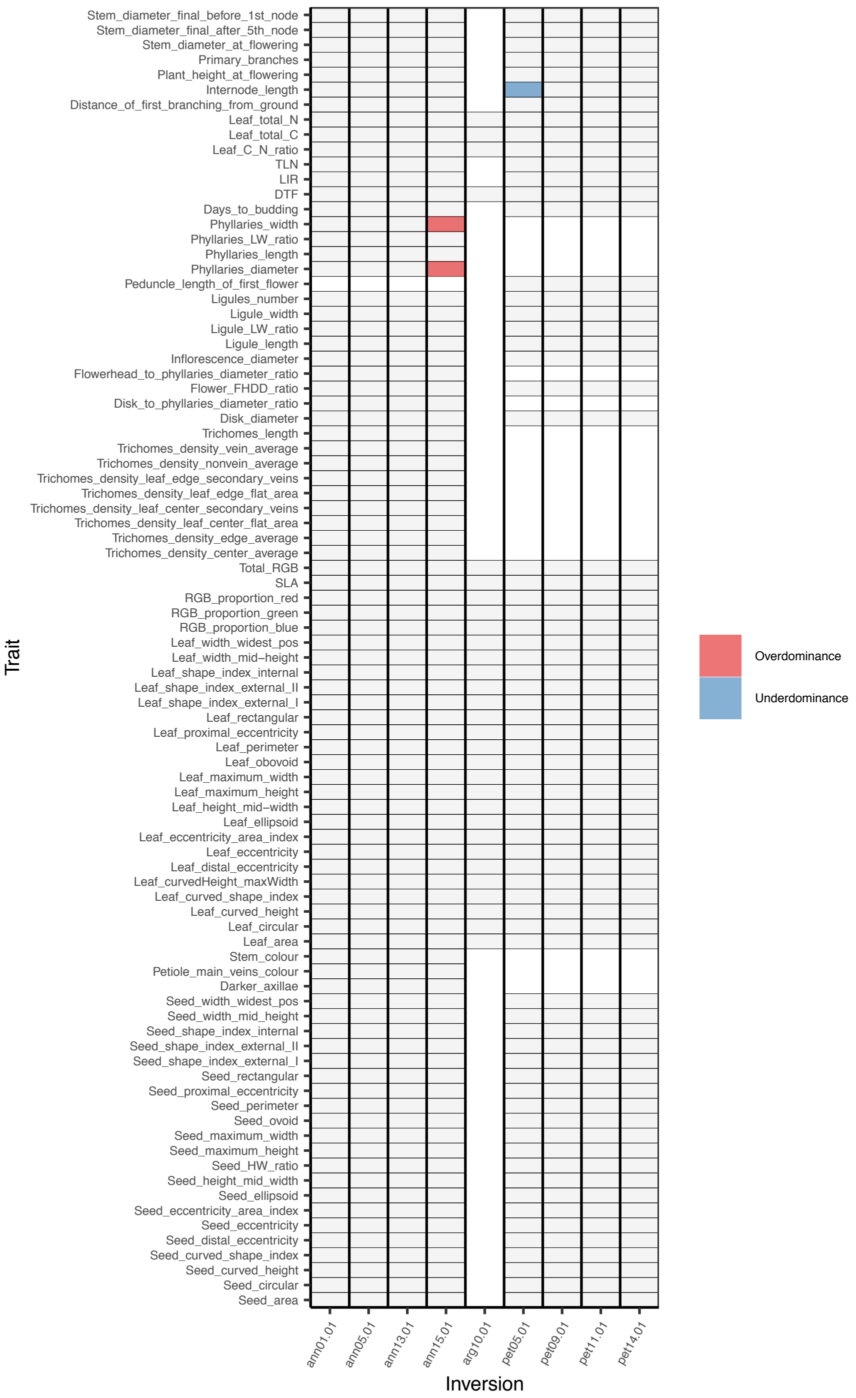


**Figure S13.** Survey of phenotypic traits across inversion genotypes. Traits surveyed for each inversion are labeled in grey. The cases where heterozygotes have a significantly ($p < 0.01$)

greater or lower phenotype than both homozygotes are shaped in color according to their condition. Trait names and details are described in Todesco et al. (2020).

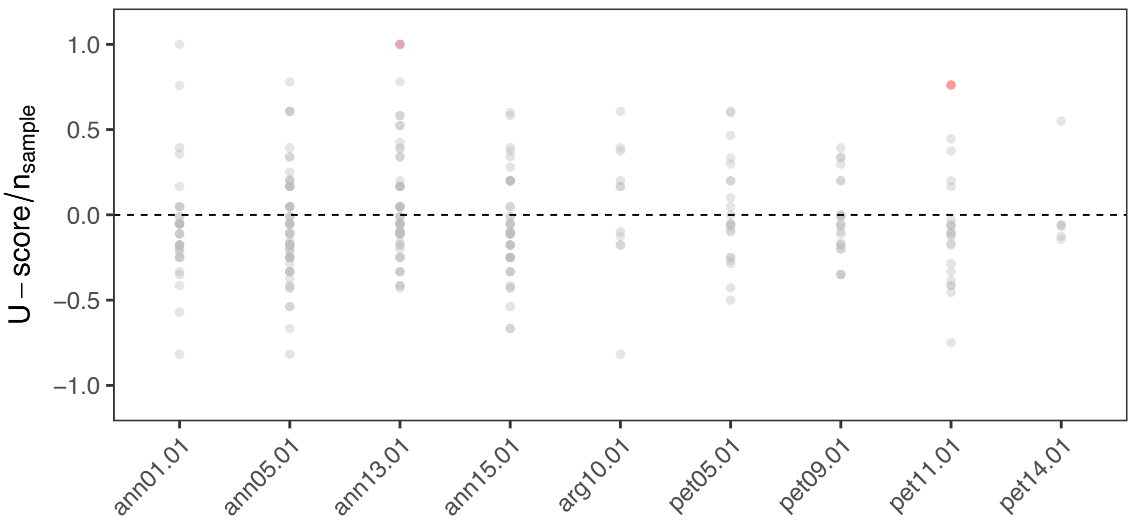


**Figure S14.** Population U-scores for inversions. U-scores were calculated for each population for each inversion and scaled by number of samples. A positive U-score indicates excess of homozygotes while a negative value indicates overrepresentation of heterozygotes. Values significantly ($p<0.01$) different from zero are marked in red.
